# Carbon dioxide shapes parasite-host interactions in a human-infective nematode

**DOI:** 10.1101/2024.03.28.587273

**Authors:** Navonil Banerjee, Spencer S. Gang, Michelle L. Castelletto, Felicitas Ruiz, Elissa A. Hallem

## Abstract

Skin-penetrating nematodes infect nearly one billion people worldwide. The developmentally arrested infective larvae (iL3s) seek out hosts, invade hosts via skin penetration, and resume development inside the host in a process called activation. Activated infective larvae (iL3as) traverse the host body, ending up as parasitic adults in the small intestine. Skin-penetrating nematodes respond to many chemosensory cues, but how chemosensation contributes to host seeking, intra-host development, and intra-host navigation – three crucial steps of the parasite-host interaction – remains poorly understood. Here, we investigate the role of carbon dioxide (CO_2_) in promoting parasite-host interactions in the human-infective threadworm *Strongyloides stercoralis*. We show that *S. stercoralis* exhibits life-stage-specific preferences for CO_2_: iL3s are repelled, non-infective larvae and adults are neutral, and iL3as are attracted. CO_2_ repulsion in iL3s may prime them for host seeking by stimulating dispersal from host feces, while CO_2_ attraction in iL3as may direct worms toward high-CO_2_ areas of the body such as the lungs and intestine. We also identify sensory neurons that detect CO_2_; these neurons are depolarized by CO_2_ in iL3s and iL3as. In addition, we demonstrate that the receptor guanylate cyclase *Ss*-GCY-9 is expressed specifically in CO_2_-sensing neurons and is required for CO_2_-evoked behavior*. Ss*-GCY-9 also promotes activation, indicating that a single receptor can mediate both behavioral and physiological responses to CO_2_. Our results illuminate chemosensory mechanisms that shape the interaction between parasitic nematodes and their human hosts and may aid in the design of novel anthelmintics that target the CO_2_-sensing pathway.

## Introduction

Skin-penetrating nematodes, including hookworms and the threadworm *Strongyloides stercoralis*, are gastrointestinal parasites that infect nearly a billion people worldwide and are agents of some of the world’s most neglected chronic diseases [1–4]. *S. stercoralis* alone infects approximately 610 million people worldwide [3]. Infections with these parasites can cause chronic gastrointestinal distress as well as stunted growth and long-term cognitive impairment in children [5–8]. Moreover, infections with *S. stercoralis* are often fatal in immunocompromised individuals [1, 5]. Current drugs used to treat these infections cannot prevent reinfection, resulting in high reinfection rates in endemic areas [1, 8]. Drug resistance due to mass drug administration is also a growing concern [9–11]. Despite the extensive morbidity caused by skin-penetrating nematodes, the basic biology of many parasitic nematodes remains incompletely understood, largely due to the challenges of working with these worms in laboratory settings. A better understanding of how skin-penetrating nematodes interact with human hosts may lead to the identification of novel drug targets that could be leveraged for nematode control.

The developmentally arrested infective third-stage larvae (iL3s) of these nematodes actively seek out hosts while in the soil environment, invade the host through skin penetration, resume development inside the host (a process called activation), and migrate within the host body to ultimately end up as parasitic adults in the host small intestine [12–14]. Host seeking, activation, and intra-host navigation – three crucial steps of the parasite-host interaction – may occur in response to host-associated chemosensory cues, yet the mechanisms that drive these processes remain obscure [15, 16]. Carbon dioxide (CO_2_) is a critical host-associated chemosensory cue for many parasites and disease vectors, including skin-penetrating nematodes [17]. Skin-penetrating nematodes encounter high CO_2_ concentrations throughout their life cycle, both in fecal and soil microenvironments and inside the host body [18–23]. Whether the high CO_2_ levels in their extra-host and intra-host microenvironments sculpt their interactions with human hosts remains poorly understood. In addition, the sensory mechanisms that drive CO_2_ responses in these nematodes are unexplored.

Here, we identify the neuronal and molecular mechanisms that drive CO_2_-evoked behavioral and physiological responses in *S. stercoralis.* We focus on *S. stercoralis* because it is unique among human-parasitic nematodes in its genetic tractability [15]. We show that *S. stercoralis* displays life-stage-specific behavioral responses to CO_2_ such that CO_2_ can be either attractive, repulsive, or neutral for *S. stercoralis* depending on the life stage. Notably, iL3s and activated iL3s (iL3as) display opposite responses to CO_2_: iL3s are repelled by CO_2_, while iL3as are attracted to CO_2_. We identify the sensory neurons (*Ss*-BAG neurons) that detect CO_2_ and promote CO_2_-evoked behavioral responses. The *Ss*-BAG neurons of iL3as display an increased CO_2_-evoked calcium response compared to iL3s, suggesting enhanced sensitivity to CO_2_ in an intra-host life stage. At the molecular level, we show that CO_2_-evoked behavioral responses are mediated by the receptor guanylate cyclase GCY-9, which is expressed specifically in the *Ss*-BAG neurons. *Ss*-GCY-9 also promotes activation of iL3s, suggesting that CO_2_ sensing via *Ss*-GCY-9 plays a crucial role in intra-host development. Drugs targeting the CO_2_-sensing pathway may thus offer a novel means of combating nematode infections. Our work illustrates how a single chemosensory cue can shape the complex interactions between parasitic nematodes and their human hosts.

## Results

### *S. stercoralis* exhibits life-stage-specific behavioral responses to CO_2_

*S. stercoralis* has a complex life cycle consisting of both extra-host and intra-host life stages [12] (Figure 1A). Parasitic adults are found in the host small intestine, where they reproduce to generate post-parasitic young larvae (L1/L2s) that are expelled in feces. Some of these larvae develop directly into iL3s; the rest undergo a single free-living generation and develop into free-living adults. The free-living adults reproduce to generate progeny that develop exclusively into iL3s. The iL3s then disperse from feces into the soil in search of hosts. Once iL3s locate a host, they invade via skin penetration and activate inside the host. The iL3as then travel through the venous bloodstream to the lungs, where they are coughed up and swallowed, ultimately ending up as parasitic adults in the small intestine [12]. However, in some cases the worms navigate directly to the small intestine without traveling through the lungs [24].

**Figure 1.**
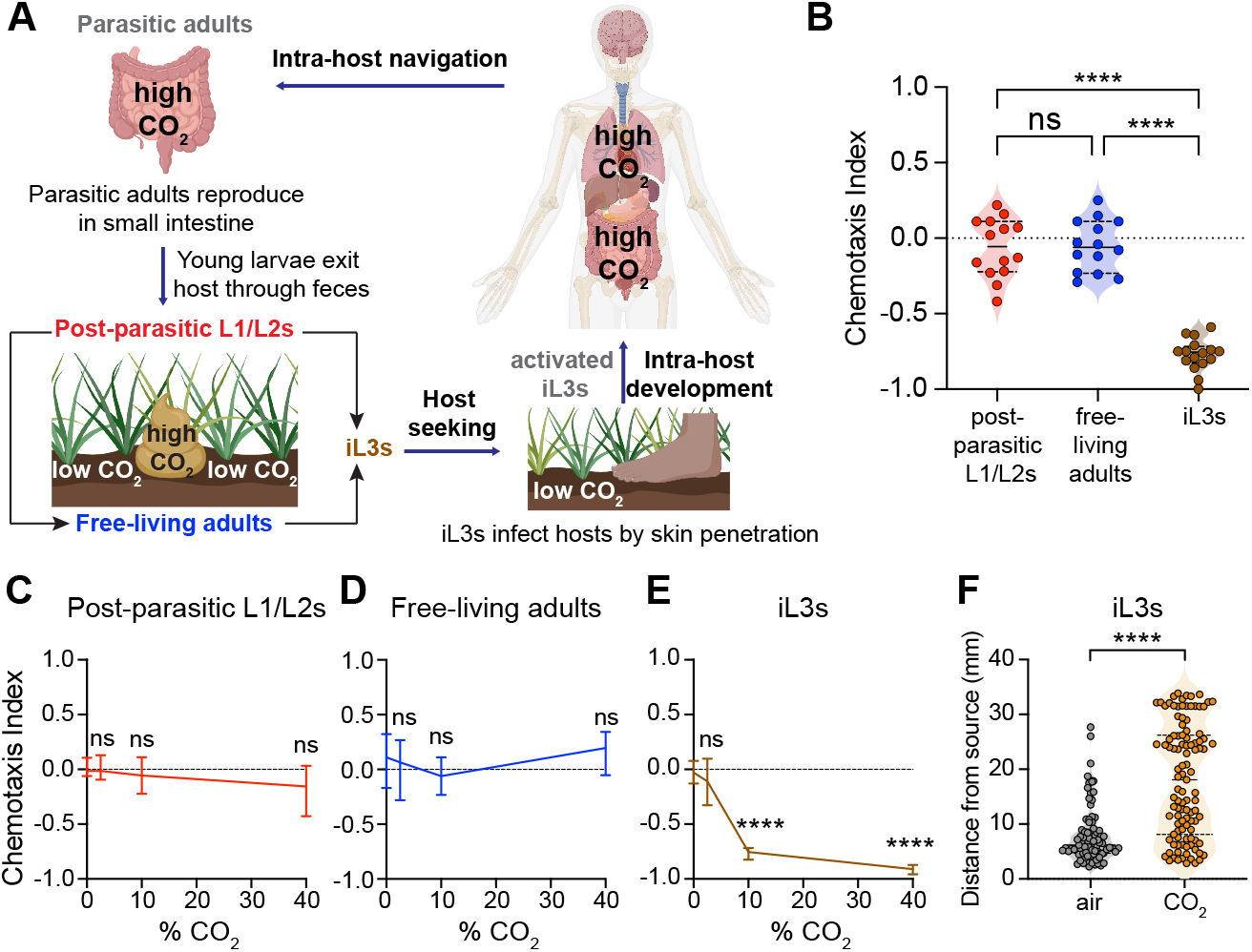
*S. stercoralis* shows life-stage-specific behavioral responses to CO_2_. **A.** The life cycle of *S. stercorali*s [12]. The progeny of parasitic adults exit the host through feces as first-stage (L1) larvae. A subset of these larvae develops directly into developmentally arrested infective third-stage larvae (iL3s), while the remaining subset cycles through a single free-living generation. All progeny of the free-living adults develop into iL3s, which seek out and invade a new host by penetrating through the skin. Upon host entry, iL3s resume development and migrate inside the host, ultimately ending up as parasitic adults in the small intestine [12]. *S. stercoralis* encounters high CO_2_ levels in host feces, host lungs, and the host small intestine [19, 20, 22, 23]. **B.** Behavioral responses of distinct life stages of *S. stercoralis* to CO_2_ in a chemotaxis assay. See also Figure S1A. The post-parasitic L1/L2 larvae and free-living adults are neutral to CO_2_, whereas iL3s are repelled by CO_2_. Each data point represents a single chemotaxis assay. n = 14-16 trials per life stage. Solid lines in violin plots show medians and dotted lines show interquartile ranges. \*\*\*\**p*<0.0001, ns = not significant, one-way ANOVA with Dunnett’s post-test. Responses are to 10% CO_2_. **C-E.** Behavioral responses of L1/L2 larvae (**C**), free-living adults (**D**), and iL3s (**E**) to CO_2_ across concentrations in a chemotaxis assay. Graphs show medians and interquartile ranges. n = 8-18 trials per life stage and condition. \*\*\*\**p*<0.0001, ns = not significant, two-way ANOVA with Dunnett’s post-test. **F.** iL3s travel significantly greater distances away from a CO_2_ source relative to an air control source in a CO_2_ dispersal assay. Each data point represents the position of a single iL3. \*\*\*\**p*<0.0001, Mann-Whitney test. n = 85-107 animals per condition. See also Figure S1B.

*S. stercoralis* encounters widely varying levels of CO_2_ throughout its life cycle that are often higher than atmospheric CO_2_ levels (∼0.04%) [18] (Figure 1A). For example, the extra-host life stages of *S. stercoralis* experience increased CO_2_ levels emitted from aerobic respiration of fecal bacteria in host feces [19, 20]. CO_2_ levels in the soil also vary from ∼0.04-13%, depending on soil depth [21]. Finally, the intra-host life stages, such as iL3as and parasitic adults, encounter elevated CO_2_ concentrations in the venous bloodstream, lungs, and small intestine; notably, CO_2_ levels can be as high as ∼35% in the small intestine [22, 23] (Figure 1A). We therefore hypothesized that CO_2_ is a critical sensory cue for *S. stercoralis* during multiple steps of its life cycle.

We first examined the behavioral responses of the environmental life stages of *S. stercoralis* to CO_2_ using a chemotaxis assay [25] (Figure S1A). We found that post-parasitic young larvae (L1/L2s) and free-living adults are neutral to CO_2_ across concentrations (Figure 1B-D). In contrast, iL3s are repelled by CO_2_ across concentrations (Figure 1B, E) [25]. What is the ecological relevance of these life-stage-specific responses to CO_2_? We hypothesized that the neutral responses of the post-parasitic L1/L2 larvae and free-living adults to CO_2_ may help them remain within host feces to feed and reproduce. In contrast, CO_2_ repulsion by iL3s may drive them to migrate off host feces and into the soil, where they can initiate host seeking. To test the hypothesis that CO_2_ acts as a dispersal cue for iL3s, we performed a CO_2_ dispersal assay in which we directly exposed populations of iL3s to a point source of high CO_2_ and then video-recorded their movement (Figure S1B). We found that iL3s traveled significantly greater distances from the CO_2_ source than from an air control (Figure 1F). These results suggest that CO_2_ avoidance may enable iL3s to migrate away from feces and into the soil to host seek.

### The receptor guanylate cyclase GCY-9 mediates behavioral and physiological responses of *S. stercoralis* to CO_2_

The receptor guanylate cyclase (rGC) GCY-9 promotes behavioral responses to CO_2_ in the free-living nematode *Caenorhabditis elegans* and is thought to function directly as a receptor for molecular CO_2_ [26, 27]. We previously identified putative rGC gene homologs in the *S. stercoralis* genome [28]. Notably, we found an *S. stercoralis* putative rGC gene homolog (SSTP_0001252500) that encodes a protein with one-to-one homology to *C. elegans* GCY-9 [28]. We named this gene *Ss*-*gcy-9*. The *Ss-gcy-9* gene encodes a protein with a domain structure resembling that of other rGC proteins – it includes an extracellular ligand-binding domain, a transmembrane domain, an intracellular kinase domain, and a guanylate cyclase domain (Figure S2A-B). To better characterize the *Ss*-*gcy-9* gene, we first analyzed its expression levels across life stages using the available life-stage-specific transcriptomic dataset for *S. stercoralis* [29–31]. We found that this gene is expressed at low levels in post-parasitic L1/L2 larvae and free-living adults but is highly expressed in iL3s (Figure 2A). The upregulation of the putative CO_2_ receptor gene *Ss*-*gcy-9* specifically in iL3s is consistent with our behavioral data showing that iL3s respond to CO_2_ (*i.e.,* they are repelled by CO_2_), whereas L1/L2 larvae and free-living adults do not respond to CO_2_ (Figure 1B-E). Taken together, these results are consistent with the possibility that GCY-9 mediates CO_2_ response and high *gcy-9* expression permits robust CO_2_-evoked behaviors in iL3s.

**Figure 2.**
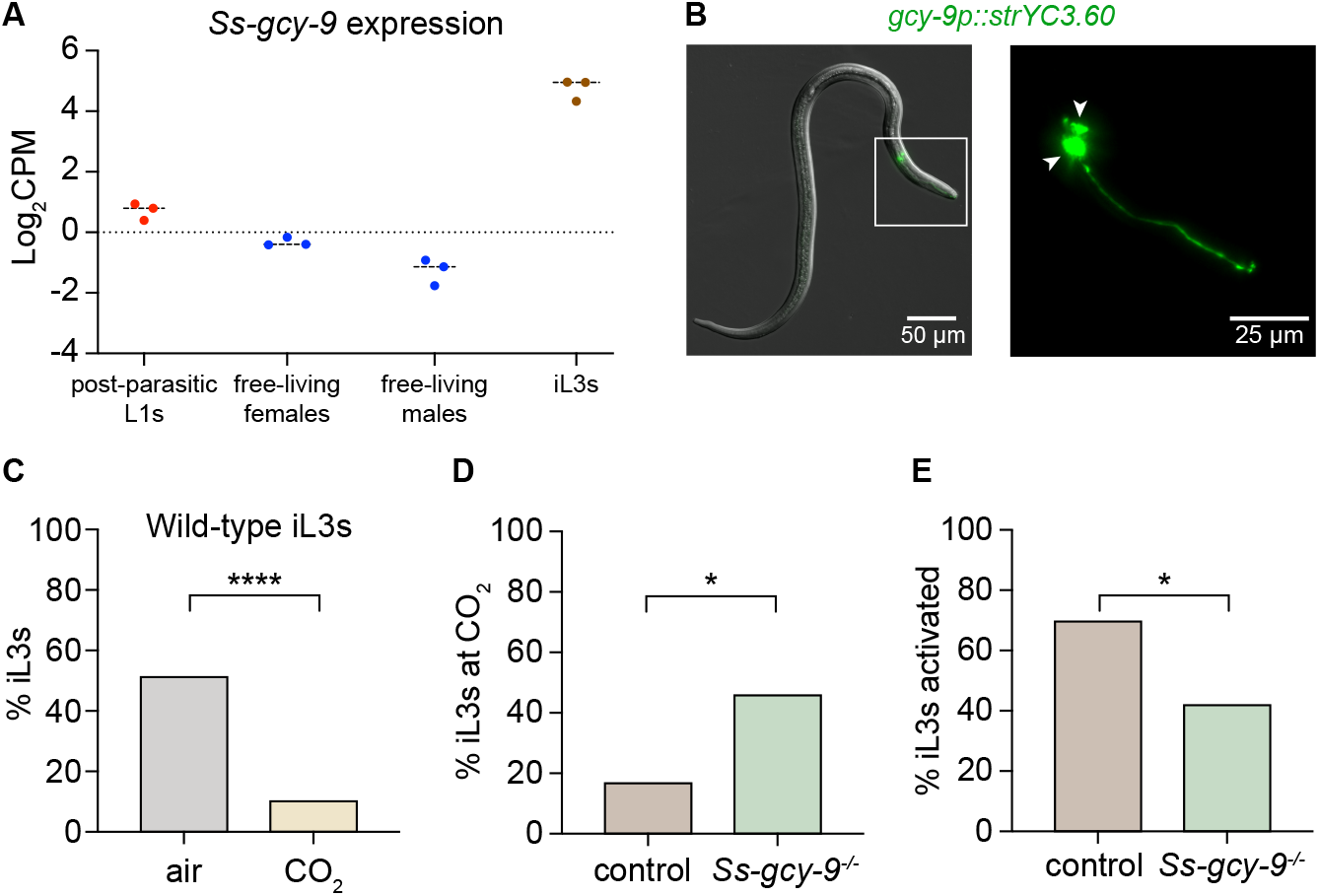
The receptor guanylate cyclase *Ss*-GCY-9 promotes CO_2_-evoked behavioral responses and activation of *S. stercoralis* iL3s. **A.** The *Ss-gcy-9* gene is transcriptionally upregulated in iL3s. Expression levels of the *Ss*-*gcy-9* gene across life stages based on whole-animal RNA-seq data [29–31]. Each data point represents a single biological replicate; lines indicate medians. Data were analyzed using the *Strongyloides* RNA-seq browser [29]. **B.** The *Ss-gcy-9* gene is expressed in the paired BAG neurons in the head. Left, representative DIC/epifluorescence overlay image of an *S. stercoralis* iL3 showing the BAG sensory neurons (green). The head is within the white box; this boxed region is enlarged in the right panel. Right, enlarged epifluorescence image of the BAG neurons. Arrowheads indicate the cell bodies; the tip of the nose is to the right. In this image, the expression of the yellow cameleon gene *strYC3.60* is driven by the *Sr-gcy-9* promoter. **C.** Behavioral responses of wild-type iL3s in single-worm CO_2_ chemotaxis assays. In the presence of a CO_2_ gradient (“CO_2_”), only ∼10% of the iL3s remain on the CO_2_ side of the plate at the end of the assay. In the absence of a CO_2_ gradient (“air”), ∼50% of iL3s remain on the CO_2_ side of the plate at the end of the assay. \*\*\*\**p*<0.0001, Fisher’s exact test. n = 69-84 iL3s per condition. **D.** *Ss-gcy-9^-/-^* iL3s are not repelled by CO_2_. Responses of no-Cas9-control and *Ss-gcy-9^-/-^* iL3s in a single-worm CO_2_ chemotaxis assay. \**p*<0.05, Fisher’s exact test. n = 24-36 iL3s per genotype. **E.** *Ss-gcy-9* contributes to activation of iL3s. Percentage of no-Cas9-control and *Ss-gcy-9^-/-^* iL3s that activated in an *in vitro* activation assay. \**p*<0.05, Fisher’s exact test. n = 31-46 iL3s per genotype. Note that in C-E, no error bars are shown because worms were scored individually; graphs display the percentage of iL3s across all assays. See also Figure S2.

We then examined the spatial expression pattern of *gcy-9*. We designed a transcriptional reporter that expresses a fluorescent marker under the control of ∼3 kb of the *Strongyloides gcy-9* promoter. We found that *gcy-9* is expressed specifically in a pair of neurons in the head region of *S. stercoralis* iL3s (Figure 2B). This expression pattern resembles that of the *C. elegans gcy-9* gene, which shows specific expression in the paired BAG sensory neurons in the head [26]. Based on conserved positional anatomy with the *C. elegans* BAG neurons and expression of the *Ss-gcy-9* ortholog, we named the GCY-9+ head neurons in *S. stercoralis* the *Ss*-BAG neurons.

To investigate whether *Ss-gcy-9* is required for the behavioral response of *S. stercoralis* iL3s to CO_2_, we disrupted the gene using CRISPR/Cas9-mediated mutagenesis (Figure S2A, C-D) [32]. We microinjected a mixture of three plasmids into the gonads of free-living adult females: 1) the expression vector for Cas9; 2) the plasmid that expresses the single guide RNA, including the target sequence specific for the *Ss-gcy-9* gene; and 3) the plasmid that supplies the repair template for inactivation of *Ss-gcy-9* by homology-directed repair. We then collected transgenic F_1_ iL3s and performed behavioral assays followed by *post hoc* genotyping to detect homozygous disruption of *Ss-gcy-9*. As a negative control, we collected F_1_ iL3s from free-living adult females microinjected with an injection mix lacking the Cas9-encoding plasmid (*i.e.*, “no-Cas9 control” iL3s). To examine the effect of *Ss-gcy-9* disruption on CO_2_-evoked behavior, we conducted single-worm CO_2_ chemotaxis assays in which we allowed the iL3 to navigate on a plate in a CO_2_ gradient for 10 min and then scored whether the iL3 was on the CO_2_ vs. air side of the plate (Figure S2E). In control assays, we allowed the iL3 to navigate on a plate for 10 min in the absence of a CO_2_ gradient but with air flowing.

In the absence of a CO_2_ gradient, ∼50% of wild-type iL3s navigated to the left side of the plate and ∼50% navigated to the right side, consistent with a random distribution across the plate. When individual wild-type iL3s were exposed to a CO_2_ gradient, only ∼10% of the iL3s remained on the CO_2_ side of the plate, consistent with wild-type iL3s being repelled by CO_2_ (Figure 2C). Like wild-type iL3s, only ∼16% of no-Cas9-control iL3s remained on the CO_2_ side of the plate. In contrast, when *Ss-gcy-9^-/-^* iL3s were exposed to the same CO_2_ gradient, ∼45% of the iL3s remained on the CO_2_ side of the plate, consistent with an almost random distribution across the plate and an inability to detect the CO_2_ gradient (Figure 2D). These results demonstrate that *Ss-gcy-9* is required for CO_2_ avoidance by *S. stercoralis* iL3s and suggest that GCY-9 plays a conserved role in CO_2_ detection in free-living and parasitic nematodes.

We then tested whether *Ss-gcy-9* is required for activation, the process whereby the developmentally arrested iL3s resume development upon host entry [33, 34]. We previously showed that CO_2_ is required to promote activation of *S. stercoralis* iL3s in an *in vitro* activation assay [35]. To test whether *Ss-gcy-9* is required for activation, we performed *in vitro* activation assays with single iL3s; in these assays, iL3s are exposed to host-like conditions and iL3as are identified based on the resumption of feeding that occurs during activation (Figure S2F) [35]. We found that significantly fewer *Ss-gcy-9^-/-^* iL3s activated, as compared with no-Cas9-control iL3s (Figure 2E). Specifically, ∼70% of no-Cas9-control iL3s vs. ∼42% of *Ss-gcy-9^-/-^* iL3s activated under similar conditions. These results suggest that *Ss-gcy-9* contributes to intra-host development in *S. stercoralis*. Together, our results identify *Ss*-GCY-9 as a putative CO_2_ receptor that drives CO_2_-evoked behavioral and physiological responses in *S. stercoralis*.

### The *Ss*-BAG sensory neurons detect CO_2_ and drive CO_2_-evoked responses

We next investigated the neural mechanisms that drive CO_2_-evoked responses in *S. stercoralis*. We focused on the *Ss*-BAG neurons, where *Ss-gcy-9* is specifically expressed (Figure 2B). To test whether these neurons detect CO_2_, we generated transgenic iL3s expressing the genetically encoded calcium indicator yellow cameleon YC3.60 specifically in the *Ss*-BAG neurons (Figure 2B). We found that the *Ss*-BAG neurons of iL3s show robust excitatory calcium responses to CO_2_ (Figure 3A-B). These responses were absent in iL3s exposed to an air control of equivalent duration (Figure S3A-B). Thus, the *Ss*-BAG neurons are activated by CO_2_ in *S. stercoralis* iL3s.

**Figure 3.**
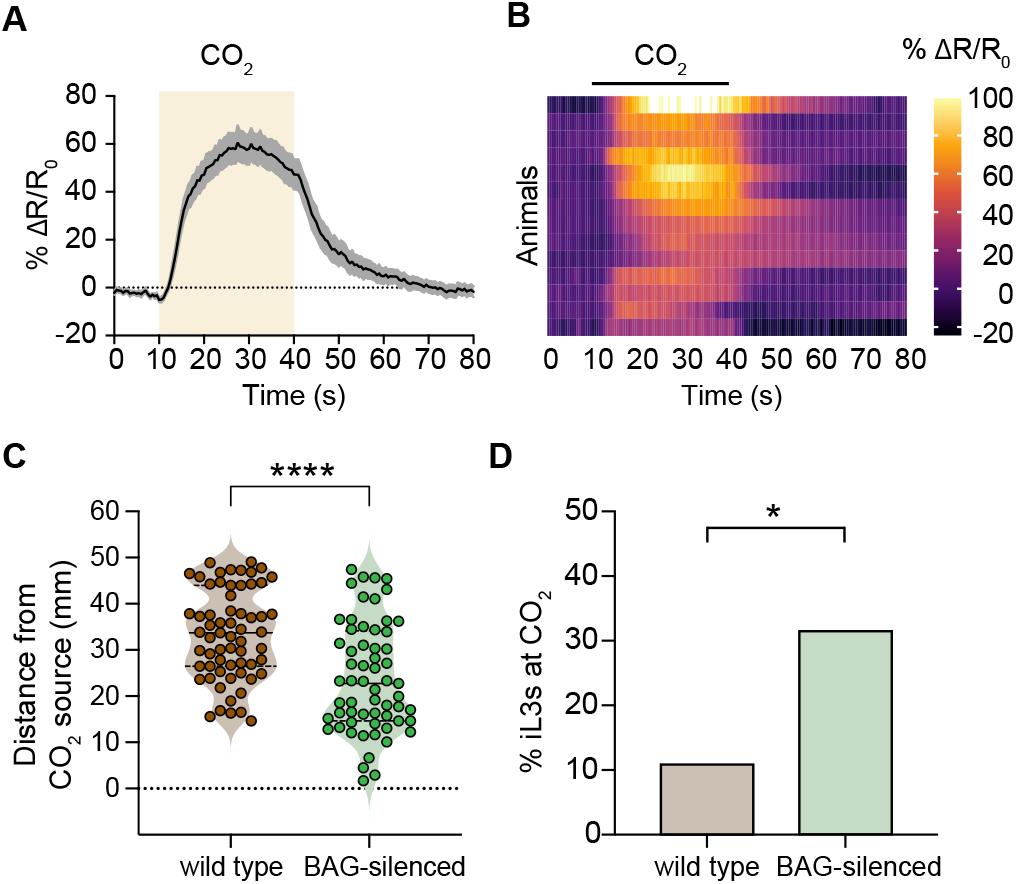
The *S. stercoralis* BAG neurons are activated by CO_2_ and mediate CO_2_-evoked behavior. **A.** *Ss*-BAG neurons are activated by CO_2_. Graph shows the calcium response (mean ± SEM) of the *Ss*-BAG neurons to a 30 s pulse of 15% CO_2_. Calcium response was measured using the ratiometric calcium indicator yellow cameleon YC3.60. The shaded box indicates the timing and duration of the CO_2_ pulse. n = 14 iL3s. **B.** Heatmap of the *Ss*-BAG calcium responses in A. Each row shows the response of a single animal. Response magnitudes (% ΔR/R_0_) are color-coded according to the scale shown to the right; rows are ordered by hierarchical cluster analysis. Black bar shows the timing and duration of the CO_2_ pulse. **C-D.** *Ss*-BAG neurons are required for CO_2_ repulsion by iL3s. Graphs show the responses of wild-type and *gcy-9p::TeTx* (“BAG-silenced”) iL3s in a CO_2_ chemotaxis assay. BAG-silenced iL3s navigate closer to the CO_2_ source (**C**) and more frequently end up on the CO_2_ side of the plate (**D**) than wild-type iL3s. \*\*\*\**p*<0.0001, Mann-Whitney test, n = 61-63 iL3s per genotype (**C**) and \**p*<0.05, Fisher’s exact test, n = 35-46 iL3s per genotype (**D**). Data shown in C-D were obtained from the same set of assays. Note that in D, no error bars are shown because the graph displays the percentage of iL3s across all assays. See also Figure S3.

To determine whether the *Ss*-BAG neurons are required to promote CO_2_-evoked behavioral responses in iL3s, we genetically silenced them by BAG-specific expression of the tetanus toxin light chain (TeTx), which inhibits neurotransmission by cleaving the synaptic vesicle fusion protein synaptobrevin (Figure S3C-E) [36, 37]. We then compared the behavioral responses of wild-type and BAG-silenced iL3s in a CO_2_ chemotaxis assay. We found that wild-type iL3s ended up significantly farther from the CO_2_ source than BAG-silenced iL3s, suggesting that BAG-silenced iL3s are less repelled by CO_2_ than wild-type iL3s (Figure 3C). The final distribution of iL3s on the plate also differed for wild-type vs. BAG-silenced iL3s, with more BAG-silenced iL3s (∼31%) ending up on the CO_2_ side of the plate than wild-type iL3s (∼11%) (Figure 3D). These results suggest that the *Ss*-BAG neurons promote CO_2_ repulsion in *S. stercoralis* iL3s. Together, our results demonstrate a crucial role for the BAG sensory neurons in detecting CO_2_ and driving CO_2_-evoked behavior in *S. stercoralis*. In addition, our results demonstrate that TeTx can be used for neuronal silencing in *S. stercoralis* as an alternative to histamine-mediated silencing using ectopic expression of the histamine-gated chloride channel HisCl1 [28, 38]. TeTx-mediated neuronal silencing is a particularly helpful addition to the parasitic nematode molecular toolkit, as it allows for interrogation of neuron function in cases where delivery of exogenous histamine is not feasible.

### *S. stercoralis* iL3as are attracted to CO_2_

Do the intra-host life stages of *S. stercoralis* also respond to CO_2_? To address this question, we focused on iL3as, the first life stage generated inside a host when iL3s exit developmental arrest and resume feeding following host invasion [12]. We activated iL3s *in vitro* (Figure S2F) and then compared the behavioral responses of iL3s and iL3as in CO_2_ chemotaxis assays performed at 37°C (to mimic host body temperature). In contrast to iL3s, which were repelled by CO_2_, iL3as were attracted to CO_2_ across concentrations (Figure 4A, S4A). To test whether the shift from CO_2_ repulsion in iL3s to CO_2_ attraction in iL3as is driven by the host-like conditions that iL3s experience during *in vitro* activation (*i.e.,* exposure to 37°C and 5% CO_2_), we performed CO_2_ chemotaxis assays with iL3s that were subjected to the activation assay but did not activate (non-activated iL3s). We found that non-activated iL3s were repelled by CO_2_ (Figure S4B). Thus, *S. stercoralis* iL3s undergo a life-stage-dependent switch in CO_2_ preference when they activate.

**Figure 4.**
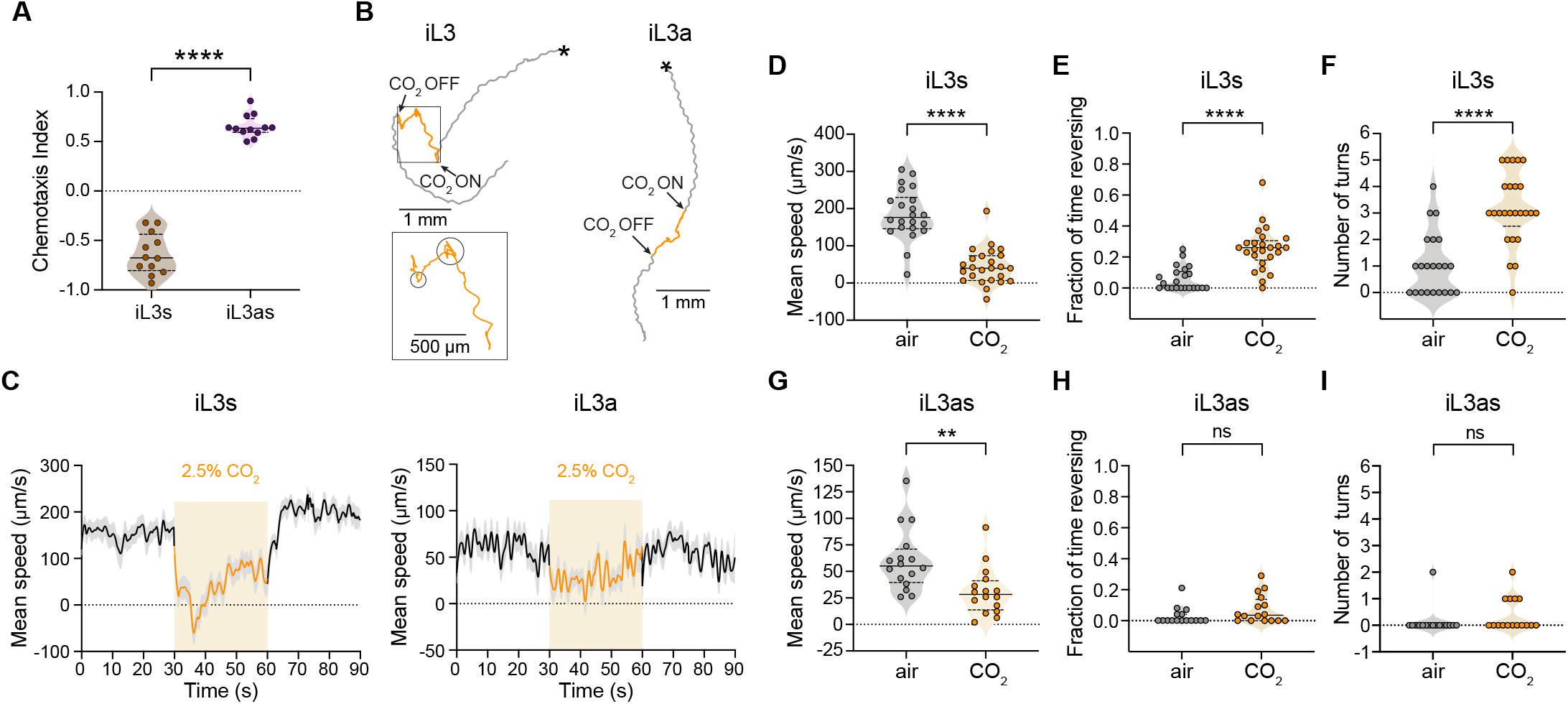
iL3as are attracted to CO_2_. **A.** iL3s are repelled by CO_2_ and iL3as are attracted to CO_2_ in a CO_2_ chemotaxis assay. Assays were conducted at 37°C to mimic the temperature of the intra-host environment; the CO_2_ stimulus contained 40% CO_2_. Each data point represents a single chemotaxis assay. Solid lines in violin plots show medians and dotted lines show interquartile ranges. \*\*\*\**p*<0.0001, Welch’s t-test. n = 12 trials per life stage. **B-I.** iL3s and iL3as respond differently to an acute CO_2_ pulse. Animals were exposed to 30 s of air, followed by 30 s of 2.5% CO_2_, followed by 30 s of air. **B.** Representative movement trajectories of an iL3 (left) and an iL3a (right) in response to a CO_2_ pulse. Asterisks indicate the position of the worm at the start of video-recording. The timing and duration of the CO_2_ pulse is indicated in orange. Enlarged region shows the movements of the iL3 during the CO_2_ pulse; regions characterized by CO_2_-evoked turns are circled. **C.** Mean smoothed instantaneous speeds (± SEM) of iL3s (left) and iL3as (right) in response to CO_2_. Shaded boxes represent the timing and duration of the CO_2_ pulse. Negative speed values indicate reverse movement. **D-F.** iL3s dramatically reduce their speed (**D**) and show an increase in reversals (**E**) and turns (**F**) in response to the CO_2_ pulse. \*\*\*\**p*<0.0001, Mann-Whitney test (**D-E**) or Welch’s t-test (**F**). n = 22-25 animals per condition. **G-I.** iL3as reduce their speed (**G**) but do not show an increase in reversals (**H**) or turns (**I**) in response to CO_2_. \*\**p*<0.01, ns = not significant, Mann-Whitney test. n = 16 animals per condition. For D-I, each data point represents the response of a single animal to a 30 s pulse of either CO_2_ or air; separate animals were exposed to CO_2_ vs. air. Solid lines in violin plots show medians and dotted lines show interquartile ranges. See also Figure S4.

We then asked whether the opposite CO_2_ preferences of iL3s (repulsion) and iL3as (attraction) are reflected in CO_2_-evoked changes in their movement patterns in response to an acute increase in CO_2_ concentration. To address this question, we video-recorded the movement patterns of animals exposed to an acute CO_2_ pulse, tracked their movement trajectories, and analyzed specific movement parameters (Figure S4C-D) [39]. We found that iL3s reduced their speed, initiated reverse movement, and increased their turn frequency in response to an acute CO_2_ pulse relative to an air control (Figure 4B-F, S4E-F), consistent with iL3s being repelled by CO_2_. In contrast, iL3as reduced their speed but did not initiate reversals and did not increase their turn frequency in response to CO_2_ (Figure 4B-C and G-I, S4E), consistent with iL3as being attracted to CO_2_. Thus, the opposing CO_2_ preferences of iL3s and iL3as are reflected in distinct CO_2_-evoked movement patterns.

### The BAG neurons of iL3as show increased CO_2_-evoked calcium activity

To investigate the neural mechanisms of CO_2_ detection in iL3as, we generated transgenic iL3as expressing YC3.60 in the *Ss*-BAG neurons. We first confirmed that the *gcy-9* promoter used to drive YC3.60 expression is expressed in the BAG neurons of iL3as (Figure S5A). We then monitored calcium responses in these neurons by exposing iL3as to CO_2_. We found that, like the BAG neurons of iL3s, the BAG neurons of iL3as show excitatory CO_2_-evoked calcium responses, suggesting that iL3as also detect CO_2_ via the BAG neurons (Figure 5A-D). However, the amplitudes of the BAG neuron responses were significantly higher in iL3as than iL3s that were exposed to the same CO_2_ concentration (Figure 5A-E). To determine whether the increased BAG calcium response in iL3as was driven by exposure of iL3s to the *in vitro* activation assay conditions (*i.e.,* 37°C and 5% CO_2_), we compared CO_2_-evoked calcium responses in the BAG neurons of iL3as and non-activated iL3s (*i.e.,* iL3s that experienced the same *in vitro* activation assay conditions but did not activate) (Figure S5B). We found that the BAG responses of iL3as were significantly higher in amplitude than those of non-activated iL3s (Figure S5C-E). Thus, the enhanced CO_2_-evoked calcium activity in the BAG neurons is specific to iL3as. Moreover, the increased calcium responses of the BAG neurons in iL3as are associated with CO_2_ attraction, since both iL3s and non-activated iL3s are repelled by CO_2_. These results provide direct evidence that an intra-host life stage of a parasitic nematode can detect a host-associated sensory cue. In addition, the elevated CO_2_-evoked calcium response of the *Ss*-BAG neurons in iL3as relative to iL3s suggests that iL3as are more sensitive to CO_2_ than iL3s.

**Figure 5.**
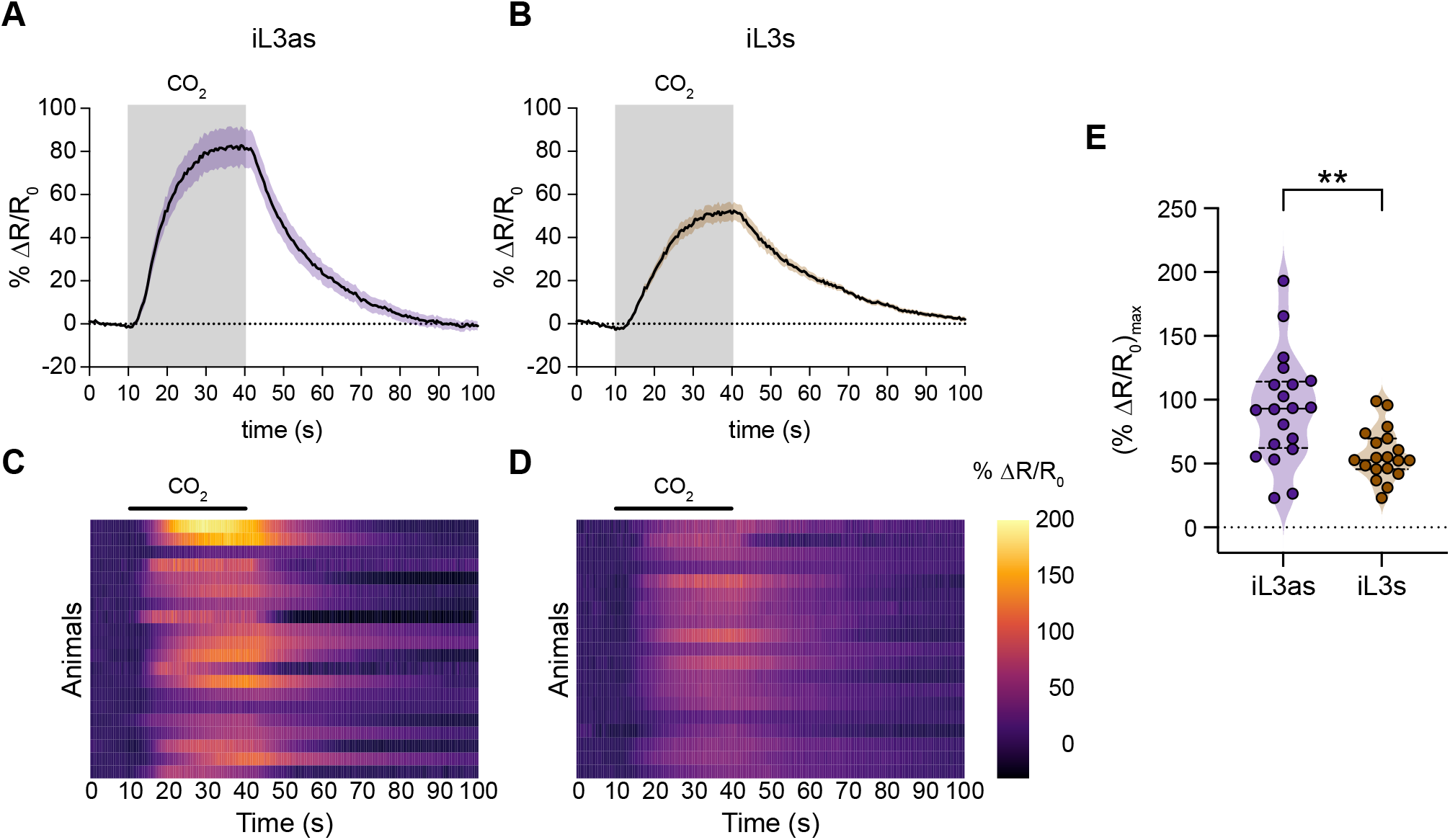
The BAG neurons of iL3as show increased CO_2_-evoked calcium responses. **A-B.** Calcium activity in the BAG neurons of iL3as (**A**) and iL3s (**B**) in response to CO_2_. Shaded boxes indicate the timing and duration of the CO_2_ pulse. Responses are to 15% CO_2_. Graphs show mean ± SEM. n = 19-20 animals per life stage. **C-D.** Heatmaps of the BAG calcium responses of iL3as (**C**) and iL3s (**D**) shown in A-B. Each row represents the response of an individual animal. Response magnitudes (% ΔR/R_0_) in the heatmaps are color-coded according to the scale shown to the right; responses are ordered by hierarchical cluster analysis. Black bars indicate the timing and duration of the CO_2_ pulse. **E.** Quantification of the maximum responses of the BAG neurons of iL3as vs. iL3s. Each data point represents the response of a single animal. Solid lines in violin plot show medians and dotted lines show interquartile ranges. \*\**p*<0.01, Welch’s *t*-test. See also Figure S5.

## Discussion

Here, we show that *S. stercoralis* exhibits robust behavioral responses to CO_2_ that vary across life stages: iL3s are repelled by CO_2_, post-parasitic young larvae and free-living adults are neutral to CO_2_, and iL3as are attracted to CO_2_. Thus, these skin-penetrating nematodes show highly dynamic responses to the CO_2_ they encounter in distinct extra-host and intra-host microenvironments throughout their life cycle. Our results support a model in which CO_2_ repulsion by iL3s drives them away from feces and into the soil environment to host seek, while CO_2_ attraction by iL3as directs intra-host migration to high CO_2_ areas of the body such as the venous bloodstream, pulmonary alveoli, and small intestine. While CO_2_ is an attractant for many parasites and disease vectors – including insect-parasitic nematodes [17], plant-parasitic nematodes [40, 41], passively ingested parasitic nematodes [42], mosquitoes [43], kissing bugs [44], and ticks [45] – the lack of CO_2_ attraction exhibited by skin-penetrating iL3s is consistent with their route of host entry: nearly all host-emitted CO_2_ is exhaled through the mouth, while only very low levels of CO_2_ escape through the skin [46–48]. However, repulsion from CO_2_ may serve a critical function in iL3s by increasing their chances of encountering a host or host-emitted attractants, such as heat or skin odorants, that direct navigation toward a host [15].

Like *S. stercoralis*, *C. elegans* exhibits CO_2_-evoked behaviors that vary across life stages. However, the homologous life stages of *S. stercoralis* and *C. elegans* exhibit very different behavioral preferences for CO_2_. In the case of *C. elegans*, well-fed adults are repelled by CO_2_ under ambient conditions [49–53]. In contrast, *S. stercoralis* free-living adults are neutral to CO_2_ (Figure 1). In addition, *C. elegans* dauer larvae and *S. stercoralis* iL3s, which are homologous life stages [54–56], show opposite responses to CO_2_; *C. elegans* dauers are attracted to CO_2_ [39, 57, 58], whereas *S. stercoralis* iL3s are repelled by CO_2_ (Figure 1) [25, 42]. *C. elegans* dauers form when environmental conditions are unfavorable, and CO_2_ attraction likely enables dauers to migrate toward potential food sources [39, 58, 59]. Thus, CO_2_ preferences of homologous life stages are not conserved across nematode species but instead reflect the ethological requirements of each species and life stage.

Previous studies of skin-penetrating nematode behavior have focused exclusively on life stages found in the environment, such as the iL3 and free-living adult stages [15, 35, 60, 61]. Although intra-host life stages have been hypothesized to use sensory cues to develop and navigate within the host body, whether these life stages exhibit sensory responses had not been tested. Here, we provide the first demonstration that an intra-host life stage exhibits a sensory-driven behavior: we show that iL3as are attracted to CO_2_ (Figure 4A). CO_2_ attraction may play an important role in directing intra-host navigation to the small intestine, where CO_2_ levels are high – either via other high-CO_2_ areas of the body such as the venous blood stream and lungs, or via direct migration to the small intestine [12, 24]. Moreover, tracking analysis revealed that iL3s and iL3as exhibit distinct movement patterns in response to an acute CO_2_ stimulus (Figure 4B-I), providing insight into their behavioral responses when they encounter abrupt changes in CO_2_ levels in their distinct microenvironments. For instance, a decrease in the speed of movement of iL3as in response to an acute increase in CO_2_ levels likely retains them in high-CO_2_ areas of the body, allowing them to complete their parasitic life cycle and successfully establish an infection. Our results illustrate the necessity of probing the sensory behaviors of the intra-host life stages of endoparasites, which may not be predictable based on the behaviors of the extra-host life stages.

As in *C. elegans*, CO_2_ response in *S. stercoralis* is mediated by GCY-9, a putative CO_2_ receptor, acting in the BAG sensory neurons; thus, conserved cellular and molecular machinery operate in free-living and parasitic nematodes at the level of CO_2_ detection (Figures 2-3) [26, 51]. In addition, our results provide insight into the sensory mechanisms that trigger activation. We show that GCY-9 drives not only CO_2_-evoked behavior but also activation (Figure 2E). Thus, CO_2_-sensing via GCY-9 mediates both behavioral and physiological responses to CO_2_. The finding that a single receptor directs both CO_2_-evoked behavior and activation raises the possibility that small-molecule inhibitors of GCY-9 could function as novel classes of anthelmintics that prevent parasitic nematodes from establishing a patent infection.

At a functional level, the BAG neurons of *S. stercoralis* iL3s are depolarized by CO_2_ (Figure 3). Moreover, the BAG neurons of iL3as show increased CO_2_-evoked calcium activity relative to the BAG neurons of iL3s (Figure 5). The increased CO_2_ sensitivity of iL3as could facilitate their retention in high-CO_2_ areas of the host body and may contribute to tissue tropism (Figure 5). We previously showed that the BAG neurons of *C. elegans* dauers are depolarized by CO_2_ [39]; thus, our results provide the first demonstration that sensory neuron response properties are in some cases conserved across free-living and parasitic nematode species. The extent to which interneuron function is conserved across species remains to be determined. In *C. elegans*, we previously showed that the CO_2_-evoked activity patterns of interneurons vary depending on the life stage of the animal [39, 58]. Thus, it is possible that the CO_2_ microcircuit that operates downstream of BAG to mediate CO_2_ response in *S. stercoralis* differs in iL3s vs. iL3as. Differences in the levels of CO_2_-evoked calcium activity in the BAG neurons of iL3s vs. iL3as may also contribute to differential activation of the downstream CO_2_ microcircuit.

Taken together, our results demonstrate that CO_2_ plays a critical role in regulating the dynamic interactions between skin-penetrating nematodes and their hosts. Our results may inform studies of CO_2_ response in other skin-penetrating, human-infective nematodes such as hookworms, which are not yet amenable to genetic manipulation. The importance of CO_2_ sensing at multiple stages of the parasite’s life cycle highlights the promise of the CO_2_-detecting pathway as a novel target for nematode control.

## Acknowledgments

We thank James Lok (University of Pennsylvania) for plasmids, Astra Bryant for generating the *Sr*-*gcy-9p::strYC3.60* construct, Manuel Zimmer (University of Vienna) for the O_2_/CO_2_ chambers used in Figures 4 and S4, and Breanna Walsh and Ruhi Patel for insightful comments on the manuscript. This work was funded by National Institutes of Health F32AI147617 to N.B., National Institutes of Health T32AI007323 (PI: P. Johnson) to N.B. and S.S.G., a UCLA Molecular Biology Institute Whitcome Fellowship to S.S.G., a UCLA Center for Academic & Research Excellence (CARE) Scholarship and National Institutes of Health R25GM055052 (PI: T. Hasson) to F.R., and National Institutes of Health R01DC021489 to E.A.H.

## Author Contributions

N.B., S.S.G., and E.A.H. conceived the study. N.B. and E.A.H. wrote the manuscript. N.B., S.S.G., M.L.C., and F.R. performed experiments and analyzed data. All authors read and approved the final manuscript.

## Methods

### RESOURCE AVAILABILITY

#### Data Availability

All data are available on GitHub at https://github.com/HallemLab/Banerjee-et-al-2024.

### EXPERIMENTAL MODEL AND SUBJECT DETAILS

All protocols and procedures involving vertebrate animals were approved by the UCLA Office of Animal Research Oversight (Protocol ARC-2011-060), which adheres to the standards of the AAALAC and the *Guide for the Care and Use of Laboratory Animals*.

#### Maintenance of *Strongyloides stercoralis*

*S. stercoralis* UPD strain was maintained in Mongolian gerbils (Charles River Laboratories) through serial passage. For infecting gerbils, *S. stercoralis* iL3s were collected from fecal-charcoal cultures using a Baermann apparatus as previously described [25, 28, 35, 62]. In some cases, iL3s were suspended in ∼0.5% low-gelling-temperature agarose and allowed to crawl out of the agarose to remove fecal debris. The iL3s that crawled out were washed 5 times with sterile 1X PBS solution. Gerbils were anaesthetized with isoflurane and inoculated with ∼2000-2250 iL3s suspended in 200 µL sterile PBS via inguinal subcutaneous injection. Following a 14-day pre-patency period, gerbil feces infested with *S. stercoralis* were collected by placing gerbils on racks overnight with wet carboard lining the cage bottom to prevent desiccation. The fecal pellets, collected the next morning, were softened with water and mixed with autoclaved charcoal granules (bone char, Ebonex Corp., Cat# EBO.5BC.04) in an approximately 1:1 ratio. The fecal-charcoal mix was stored in 10-cm diameter x 20-mm height Petri dishes lined with filter paper moistened with dH_2_O. Gerbil feces infested with *S. stercoralis* were collected from days 14-45 post-infection.

#### Generation of transgenic or mutant *S. stercoralis*

Free-living *S. stercoralis* adult females were microinjected with plasmid constructs and transgenic F_1_ iL3s were obtained as previously described [63]. Microinjected females were incubated on fecal-charcoal plates with wild-type adult males at 23°C for at least 6 days prior to screening and performing assays. A full list of plasmids used in this study is provided in Table S1. The *Ss-gcy-9* gene was disrupted using established CRISPR/Cas9 methods [28, 32, 62]; a detailed description of targeted mutagenesis of *Ss-gcy-9* is provided below. For calcium imaging, transgenic F_1_ iL3s expressing *strYC3.60* in the *Ss*-BAG neurons were generated by microinjecting the pASB55 construct at a concentration of 50 ng/µL. For silencing the *Ss*-BAG neurons, transgenic F_1_ iL3s expressing *strTeTx* in the *Ss*-BAG neurons were generated by microinjecting the pNB11 construct at a concentration of 50 ng/µL.

### METHOD DETAILS

#### Molecular biology

The promoter sequence for the *gcy-9* gene was extracted from *Strongyloides ratti* genomic DNA by PCR-amplifying ∼3 kb upstream of the predicted first exon and a portion of the first exon using primers MLC168 and MLC169 (Table S2). The putative promoter sequence was cloned into the *Strongyloides* expression vector pPV254, which contains *Ss-act-2p*::GFP [64], in place of the *Ss-act-2* promoter to generate the *Sr-gcy-9p::GFP* (pMLC29) construct. For expression of the calcium indicator yellow cameleon YC3.60 in *S. stercoralis* BAG neurons, the *Sr-gcy-9* promoter sequence from pMLC29 was subcloned into pASB52, which contains the *Strongyloides*-codon-optimized *YC3.60* sequence (*strYC3.60*) [28], to generate pASB55, the *Sr-gcy-9p::strYC3.60* construct. For expression of tetanus toxin in the *Ss*-BAG neurons, we first generated a *Strongyloides*-codon-optimized *TeTx* sequence (*strTeTx*) using the Wild Worm Codon Adapter [65]. To generate pNB11, which contains *Sr-gcy-9p::strTeTx::P2A::strmScarlet-I*, the *strTeTx::P2A::mScarlet* sequence was synthesized by GenScript (Piscataway, NJ). The *Sr-gcy-9* promoter was then subcloned from pMLC29 into the *strTeTx::P2A::mScarlet* construct to generate pNB11.

#### *In vitro* activation assays

*In vitro* activation assays with *S. stercoralis* iL3s were performed as previously described [33–35]. For the population assays shown in Figures 4A and S4A-B, iL3s were collected from a Baermann apparatus; washed 3 times in BU saline [66] in 15 mL conical tubes; and resuspended in 10 mL BU saline supplemented with 100 µL of 100x penicillin-streptomycin (10,000 u/mL, Gibco 15140-122), 100 µL of 100x amphotericin B (250 µg/mL, Gibco 15290-018), and 10 µL of 1000x tetracycline hydrochloride (5mg/mL, Sigma-Aldrich T7660-5G) dissolved in ddH_2_O. iL3s were axenized in the dark (tetracycline hydrochloride is light-sensitive) for 3 h at room temperature. iL3s were then pelleted by centrifugation and the supernatant was removed. iL3s were titrated to a concentration of ∼50 iL3s per µL of BU saline and 2-4 µL of the pelleted iL3s (containing ∼100-200 iL3s) was transferred to each well of a 96-well plate containing 110 µL DMEM (Corning 10-013-CV) supplemented with antibiotics (same final concentrations as above) that was preincubated at 37°C and 5% CO_2_ for 1-3 h. For population assays, iL3s were added to 12-36 wells containing DMEM. The 96-well plate containing iL3s was then incubated at 37°C with 5% CO_2_ for 21 h, after which 2.5 µL of fluorescein isothiocyanate dye (FITC, Acros Organics 119252500, 20 mg/mL in N,N-dimethylformamide) was added to each well. iL3s were incubated with FITC at 37°C and 5% CO_2_ for an additional 3 h. iL3s were then collected from all wells into a 15 mL conical tube filled with BU saline and washed 5 times with BU saline to remove excess FITC before transferring to chemotaxis assay plates.

For *in vitro* activation assays with single no-Cas9-control or *Ss-gcy-9^-/-^* iL3s, as shown in Figure 2E, a single iL3 was axenized in 110 µL of BU saline supplemented with antibiotics, as described above, in one well of a 96-well plate. Each iL3 was then transferred into a well containing DMEM (Corning 10-013-CV) and was incubated as described above. After incubation, each iL3 was washed to remove FITC by pipetting up and down in a watch glass with ∼2 mL dH_2_O. The iL3 was then transferred in a single droplet to a 2% Nematode Growth Media (NGM) agar plate [67] and paralyzed with nicotine (Sigma N3876, 1% in dH_2_O). The iL3 was then scored for activation based on the presence (activated) or absence (non-activated) of FITC in the pharynx, using a Leica M165 FC fluorescence microscope. Individual iL3s were then retrieved from the plate for genomic DNA isolation and genotyping, as described below. All genotypes were tested in parallel and experiments were conducted over multiple days to account for day-to-day variability in assay conditions. iL3s were tested blind to genotype and were genotyped *post hoc*.

For calcium imaging, transgenic iL3s expressing yellow cameleon YC3.60 were activated using DMEM (Gibco 11995-065) in a watch-glass under the same conditions as above, except that the red fluorescent dye Alexa Fluor NHS Ester 594 (ThermoFisher A20004, 1 mg/mL in N,N-dimethylformamide) was used, instead of FITC, as an indicator for activation. 25 µL of 1 mg/mL Alexa dye was added to 1 mL media containing iL3s. iL3as and non-activated iL3s for calcium imaging were screened and sorted based on the presence or absence of Alexa Fluor in the pharynx, respectively, using a Leica M165 FC fluorescence microscope.

### Preparation of animals for CO_2_ chemotaxis assays

For all life stages except iL3as, animals were isolated using a Baermann apparatus. Post-parasitic L1/L2 larvae and iL3s were collected in a 15 mL conical tube and free-living adults were collected in a 65 mm Syracuse watch glass. For iL3s and free-living adults, animals were washed twice with BU saline and then once with ddH_2_O in the conical tube (for iL3s) or the watch glass (for free-living adults) prior to transfer to assay plates. The post-parasitic L1/L2 larvae were washed 4 times with BU saline and then once with ddH_2_O in the conical tube prior to transfer to assay plates. *S. stercoralis* iL3s were collected from fecal-charcoal plates incubated at 23°C for 7-10 days using a Baermann apparatus set up for 30-90 min. After washing, iL3s were transferred in droplets onto assay plates. Young free-living adults (males and females) were collected from fecal-charcoal plates incubated at 25°C for 1 day using a Baermann apparatus set up for 1-2 h. After washing, adults were gently transferred onto a small rectangular piece of Whatman filter paper, which was used to transport them to assay plates. Post-parasitic L1/L2 larvae were isolated from fecal pellets collected the same day and not mixed with charcoal granules, using a Baermann apparatus set up for ∼3 h. Following washing, the L1/L2 larvae were transferred in droplets onto assay plates. iL3as were collected from *in vitro* activation assays (see above), washed 5 times with BU saline and once with ddH_2_O, and then transferred in droplets onto assay plates. For data shown in Figures 2D and 3C-D, the F_1_ iL3 progeny from microinjected adult females were collected using a Baermann apparatus (after at least 6 days post-microinjection), washed twice with BU saline, and kept in a small watch glass in BU saline. iL3s were then screened by placing ∼100-200 iL3s on a 6 cm 2% NGM plate seeded with a thick lawn of *Escherichia coli* OP50; freely-moving transgenic iL3s were picked and transferred to a small watch glass with ∼1 mL BU saline. Prior to assays, iL3s were washed once with ddH_2_O in a watch glass and transferred to assay plates in 2-3 µL droplets. No-Cas9-control iL3s and wild-type control iL3s were isolated and treated similarly; moreover, assays with mutant iL3s were performed on the same day as assays with no-Cas9-control iL3s (Figure 2D), and assays with transgenic iL3s were performed on the same day as assays with wild-type control iL3s (Figure 3C-D). Assays were conducted over multiple days to account for day-to-day variability.

### CO_2_ chemotaxis assays

CO_2_ chemotaxis assays were performed essentially as previously described [25, 42]. Animals were placed onto the center of a 9 cm 2% NGM agar plate at the start of the assay. Gases comprising a CO_2_ stimulus (the desired concentration of CO_2_, 21% O_2_, balance N_2_) and an air stimulus (21% O_2_, balance N_2_) were pumped through holes in opposite sides of the plate lid to establish a CO_2_ gradient. A syringe pump (PHD 2000, Harvard Apparatus) was used to deliver gas stimuli through ¼-inch flexible PVC tubing at flow rates of 1.5 mL/min (for iL3as) and 0.5 mL/min (for all other life stages). Assays ran for 30 min (for iL3as) or 1 h (for all other life stages). For iL3as, assays were conducted in a 37°C incubator; assay plates and gases (within syringes) were pre-incubated at 37°C for ∼30 min prior to assays. For data shown in Figure 4A, assays with iL3s were performed under the same conditions as iL3as. The number of animals that navigated into a 20 mm diameter circle under each gas inlet was counted at the end of the assay. For data shown in Figures 1B and 1D, the total number of free-living adults that navigated to either scoring region were counted; males and females were not counted separately. For data shown in Fig. 4A and Fig. S4A, only worms with a green pharynx (due to ingestion of FITC) were counted. For data shown in Figure S4B, iL3as and non-activated iL3s (as identified by the presence or absence of FITC in the pharynx, respectively) were counted separately within each scoring region of the same assay plate using a Leica M165 FC fluorescence microscope. A chemotaxis index (CI) was then calculated as:

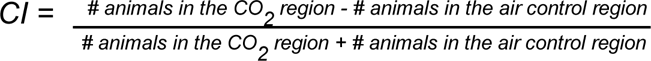

To account for directional bias due to vibration, assays were conducted in pairs, with the CO_2_ gradients in opposite orientations for the two plates. If the absolute value of the difference in CI between two assays in a pair was ≥0.9, both assays were discarded as behavior was assumed to be impacted by directional bias. Assays were also discarded if fewer than 7 animals moved to the combined scoring regions.

For data shown in Figures 2C and 3C-D, 2-10 iL3s were placed onto the center of the assay plate. For data shown in Fig. 2D, a single iL3 was placed onto the center of the assay plate. Assays ran for 10 min. For data shown in Figures 2C-D and 3D, iL3s that navigated to or remained within a 1-cm-diameter region down the center of the plate were excluded from analysis. For data shown in Figure 3C-D, the responses of iL3s were video-recorded using a Leica S9D microscope (equipped with a 0.5x supplementary lens, a 300 mm M-series Focus drive, and a TL3000 Ergo transmitted light base) with an attached Basler Ace 20-megapixel acA5472-17µm camera (mounted on a 0.63x Leica adapter lens). Image sequences were captured at 0.5 frame/s using Pylon viewer software (Basler). Image sequences were processed to generate videos using Fiji [68]. For this experiment, the gas inlets were placed closer (3.5 cm apart) for video-recording. For Figure 3C, the final distance of each iL3 was calculated from a fixed point at the CO_2_ inlet, either at the end of the assay (last captured frame) or before the iL3 left the field of view using Fiji [68]. For Figures 2D and 3D, iL3s were tested blind to genotype and genotypes were revealed *post hoc*.

### CO_2_ dispersal assays

*S. stercoralis* iL3s were isolated using a Baermann apparatus as described above, washed twice in BU saline, and stored in a watch glass containing 1 mL BU saline. CO_2_ dispersal assays were performed using 9 cm 2% NGM agar plates with a lid containing a single hole in the center for gas delivery. Prior to assays, plates were left in a fume hood with the lids off for 90 min to remove excess moisture from the agar surface, and then left on the bench with the lids on at room temperature for at least 30 min. ∼20-30 iL3s were then transferred in a 3 µL droplet to the center of the assay plate. The plate was immediately covered with the lid fitted with flexible PVC tubing (1/16-inch ID x 1/8-inch OD) through which a gas mixture – either 10% CO_2_, 21% O_2_, balance N_2_ (CO_2_ stimulus) or 21% O_2_, balance N_2_ (air control) – was delivered at a flow rate of 2 mL/min using a syringe pump (PHD 2000, Harvard Apparatus). Video-recording started immediately after the droplet dried and animals started crawling on the agar surface. Animals were video-recorded and processed as described above, except that image sequences were captured at 2 frames/s. Animals were video-recorded for 5 min. The final distance of each iL3 from the gas source was calculated at the end of the assay (last captured frame) or just before the iL3 left the field of view using Fiji [68].

### CRISPR/Cas9-mediated targeted mutagenesis of *Ss*-*gcy-9*

The *Ss-gcy-9* gene, SSTP_0001252500, was identified as an ortholog of *C. elegans gcy-9* using a BLAST search on WormBase ParaSite [28, 32, 62, 69, 70]. The CRISPR target site (AATCTTAAATCAAAAGGTGG) and guide RNA were selected using Geneious 9 software, as previously described [32, 62]. The single guide RNA (sgRNA) expression construct targeting *Ss-gcy-9* (pSSG05) was synthesized by Genewiz (South Plainfield, New Jersey) to include 500 bp of the *S. ratti* U6 promoter and 277 bp of the *S. ratti* U6 3’ UTR flanking the *Ss-gcy-9* sgRNA. For homology-directed repair at *Ss-gcy-9*, a repair construct (pSSG04) was made by subcloning 570 bp 5’ and 583 bp 3’ homology arms flanking the *Ss-gcy-9* CRISPR site into the pAJ50 construct containing an *Ss-act-2::mRFPmars* cassette, which drives *mRFPmars* expression in the body wall muscle [64]. Cas9 endonuclease was expressed from the vector pPV540, where *Strongyloides*-codon-optimized Cas9 expression is driven by the *S. ratti eef-1A* promoter [32, 71]. The pPV540, pSSG04, and pSSG05 constructs were mixed and microinjected into free-living adult females as described above. F_1_ iL3 progeny were screened for potential *Ss-gcy-9* disruptions as described below. To obtain *Ss-gcy-9^-/-^* iL3s, adults were microinjected with 80 ng/µL pSSG04, 80 ng/µL pSSG05 and 40 ng/µL pPV540 constructs. To obtain no-Cas9-control iL3s, microinjections were performed using the same recipe but with pPV540 omitted from the mix. A summary of plasmids is provided in Table S1.

The F_1_ iL3 progeny obtained from microinjected adult females were isolated using a Baermann apparatus. The iL3s were washed twice in BU saline and stored in BU saline in a watch glass. Transgenic iL3s were screened for *mRFPmars* expression by pipetting ∼15-20 µL of iL3s in BU saline (∼100-200 iL3s) onto a 6 cm 2% NGM plate seeded with *Escherichia coli* OP50. Freely moving iL3s were screened for *mRFPmars* expression using a Leica M165 FC fluorescent microscope; transgenic iL3s were picked and stored in a small watch glass with ∼1 mL of BU saline prior to assays. Only iL3s with near-uniform *mRFPmars* expression along the full body wall were picked as they were more likely to have undergone homology-directed repair; iL3s with patchy or faint *mRFPmars* expression were not collected as they were more likely to only express *mRFPmars* from extrachromosomal arrays [32]. Single-worm chemotaxis assays were performed using transgenic iL3s as described above, followed by *post-hoc* genotyping to confirm integration of the repair template at the target region. Additional experimental details can be found in Table S3.

### Single iL3 genotyping

Genomic DNA was extracted from single iL3s as previously described [28, 32, 62]. A single iL3 was collected in a PCR tube containing 5-6 µL of nematode lysis buffer (50 mM KCl, 10 mM Tris pH 8, 2.5 mM MgCl_2_, 0.45% Nonidet-P40, 0.45% Tween-20, 0.01% gelatin in dH_2_O) supplemented with ∼0.12 µg/µL Proteinase-K and ∼1.7% 2-mercaptoethanol. Tubes were kept at −80°C for at least 20 min, and then placed in a thermocycler for digestion: 65°C (2 h), 95°C (15 min), 10°C (hold). For long-term storage before digestion, iL3s were kept at −80°C and digestion was performed on the same day as PCR genotyping. To genotype iL3s, PCR reactions were performed with GoTaq G2 Flexi DNA Polymerase (Promega, Cat. #M7801) or Herculase II Fusion DNA Polymerase (Agilent, Cat. #600675) using the following thermocycler conditions: denature 95°C (2 min); PCR 95°C (20 s), 55°C (20 s), 72°C (1 min) x 35 cycles; final extension 72°C (5 min); 10°C (hold). The single iL3 digests were split evenly across the control, wild-type locus, and 5’ integration reactions, as shown in Figure S2D. Primer sets used for *Ss-gcy-9* genotyping can be found in Table S2.

### gcy-9 gene structures and GCY-9 protein domain annotations

The gene structure diagrams for *C. elegans* and *S. stercoralis gcy-9* shown in Figure S2A were generated using Exon-Intron Graphic Maker (Version 4, www.wormweb.org). The predicted protein domains of *C. elegans* and *S. stercoralis* GCY-9 shown in Figure S2B were annotated using InterPro [72]. The schematics of the GCY-9 protein domains were generated using the MyDomains: Image Creator function of PROSITE (http://prosite.expasy.org/mydomains/).

### Calcium imaging

Calcium imaging experiments were performed on animals expressing a *Strongyloides*-codon-optimized version of the genetically encoded calcium indicator gene *yellow cameleon YC3.60* [28]. For data shown in Figures 3A-B and S3A-B, transgenic *S. stercoralis* iL3s expressing YC3.60 in the *Ss*-BAG neurons were screened in 1% nicotine as described above and stored in BU saline to recover overnight prior to imaging. For data shown in Figures 5B and D, transgenic iL3s moving on an *E. coli* OP50 lawn were collected and stored in BU saline overnight prior to imaging. For data shown in Figures 5A, 5C, and S5, iL3as and non-activated iL3s were generated as described above; moving animals of each category were screened (for the presence or absence of Alexa Fluor in the pharynx) on an unseeded 2% NGM agar plate and separately collected in watch glasses with BU saline.

To immobilize animals for imaging, a 2% agarose pad (made with ddH_2_O; <10 mm in diameter) was made on a 48 x 60 mm cover glass (Brain Research Laboratories, Cat # 4860-1D) and left to dry overnight. A single animal was transferred in a droplet of BU saline onto the dry agarose pad. The droplet was then absorbed using Whatman paper, leaving the animal behind. This resulted in attachment of the animal to the dry agarose pad, which restricted its movement during imaging. A perfusion chamber of 20 mm diameter and 2.5 mm height (Grace Bio-Labs CoverWell, Millipore Sigma, Cat. #GBL622301) with an adhesive base and two 1.5 mm ports (with attached tubing connectors, Grace Bio-Labs press fit tubing connectors, Millipore Sigma, Cat. #GBL460003) on diametrically opposite sides for gas inlet and outlet, was placed on the cover glass around the animal. Humidified gas was delivered through one port into the chamber through flexible PVC tubing at a flow rate of 30 mL/min (controlled by a flow meter, VWR model #GR60140AVB-VW) from two gas tanks fitted with valves controlled by a ValveBank TTL pulse generator. An air pulse (21% O_2_, balance N_2_) was delivered for 30 s, followed by a 30 s CO_2_ pulse (15% CO_2_, 21% O_2_, balance N_2_) and then another air pulse (21% O_2_, balance N_2_) for 60 s. For air controls, the CO_2_ pulse was replaced with an air pulse of equivalent duration (30 s). Air stimuli were delivered to separate sets of worms in control experiments. To check for a complete seal of the perfusion chamber on the cover glass, flexible PVC tubing connected to the gas outlet was immersed in dH_2_O in a watch glass; water bubbles in the watch glass ensured a seal. Imaging was performed using a Zeiss AxioObserver A1 inverted microscope equipped with a 40x EC Plan-NEOFLUAR objective lens, a Colibri 7 (Zeiss) for LED fluorescence illumination, a 78 HE ms (1) filter set (BP445/25 + BP510/15, DFT460+520; Zeiss) and a Hamamatsu ORCA-Flash4.0 camera for simultaneous acquisition of CFP and YFP images. Images were acquired in the YFP and CFP channels at 2 frames/s using Zeiss ZEN 3.4 (blue edition) software. The emission image was passed through a Hamamatsu W-View Gemini beam splitter with a CFP/YFP dual camera filter set.

Image processing and analyses were performed using Zen 3.4 (blue edition) software and Microsoft Excel. Images were analyzed by selecting two regions of interest (ROIs) - one ROI consisted of the soma of the *Ss*-BAG neuron and the other ROI consisted of a background region. The average intensity for YFP and CFP of the background region was subtracted from the average intensity for YFP and CFP of the soma. YFP values were adjusted to correct for CFP signal bleed-through, and the YFP/CFP ratio was then calculated. For each dataset, the different life stages or conditions were tested in parallel across multiple days. For quantification, the response period was defined as the time interval beginning with the onset of the CO_2_ pulse and ending 10 s after the offset of the CO_2_ pulse. The % ΔR/R_0_ (max) values were calculated during the response period. Graphs and heatmaps were generated using GraphPad Prism v9.1.0. Heatmaps were generated using the web-based tool Heatmapper [73]; responses within heatmaps were ordered by hierarchical clustering analysis, using Euclidean distance as a similarity measure.

### Fluorescence microscopy

For microscopy, iL3s, iL3as, and non-activated iL3s were screened for YC3.60 expression, mScarlet-I expression, and/or for the presence of Alexa Fluor on a Leica M165 FC fluorescence microscope following nicotine paralysis [32]. Animals were collected in a watch glass with BU saline. Animals were then mounted in droplets on a slide with a 5% Noble agar (dissolved in BU saline) pad, exposed to 100 mM levamisole (dissolved in BU saline), and covered with a cover slip. Epifluorescence and DIC imaging were performed using an inverted Zeiss AxioObserver A2 microscope equipped with a Plan-APOCHROMAT 20X objective lens, a Colibri 7 (Zeiss) for LED fluorescence illumination, a 38 HE filter set for GFP (BP470/40, FT495, BP 525/50), a 63 HE filter set for mScarlet-I or Alexa Fluor (BP572/25, FT590, BP629/62), a Hamamatsu ORCA-Flash4.0 camera, and Zen 3.3 (blue edition) software (Zeiss). Images were captured as z-stacks and maximal intensity projections were constructed using Fiji [68].

### Acute CO_2_ assays and behavioral tracking

Acute CO_2_ assays were performed as previously described [39, 74], with modifications *S. stercoralis* iL3s were isolated using a Baermann apparatus, washed twice with BU saline in a 15 mL conical tube, and then kept in a watch glass with ∼1 mL BU saline. iL3as were generated as described above; moving animals were screened for the presence of FITC in their pharynx on an unseeded 2% NGM agar plate and stored in a watch glass in ∼1 mL BU saline. Both iL3s and iL3as were washed with ddH_2_O in a watch glass before transferring to assay plates.

Assays were performed on 14 cm unseeded 2% NGM plates at room temperature. Prior to assays, plates were left in a fume hood with the lids off for 1 h to remove excess moisture from the agar surface, and then on the bench (lids on) at room temperature for at least 30 min. 6-10 animals were transferred in water droplets onto assay plates, the droplets were allowed to dry, and the animals were then left to acclimate on the agar surface for ∼1 min. A chamber with a 6 cm viewing arena, connected to ¼-inch flexible PVC tubing on opposite ends for a gas inlet and outlet (Figure S4C), was placed on top of the assay plate as previously described [39, 74]. Premixed gases were delivered into the chamber through the gas inlet at a flow rate of ∼700 mL/min (controlled by a flow meter, VWR model #FR2A138VVT-VW) from gas tanks (Airgas) fitted with valves controlled by a ValveBank. Prior to video-recording, air (21% O_2_, balance N_2_) was delivered into the chamber for 1 min to acclimate animals to gas flow. Video-recording started immediately post-acclimation, when animals were exposed to a 30 s pulse of air (21% O_2_, balance N_2_), followed by a 30 s pulse of CO_2_ (2.5% CO_2_, 21% O_2_, balance N_2_) and then a 30 s air pulse (21% O_2_, balance N_2_) (Figure S4D). For air controls, the CO_2_ pulse was replaced with an air pulse of equivalent duration (30 s). To check for a complete seal of the chamber on the agar surface of the assay plate, the gas outlet was immersed in dH_2_O in a conical tube; water bubbles in the tube ensured a seal). Animals were videorecorded for a total of 90 s using a Leica S9D microscope (equipped with a 0.5x supplementary lens, a 300 mm M-series Focus drive, and a TL3000 Ergo transmitted light base) with an attached Basler Ace 20-megapixel acA5472-17µm camera (mounted on a 0.63x Leica adapter lens). Image sequences were captured at 10 frames/s using Pylon viewer software (Basler). Image sequences were processed to generate videos using Fiji [68].

Tracking and analyses of movement trajectories were performed using WormLab 2022.1.1 (MBF Bioscience LLC, Williston, VT USA). Videos were optimally thresholded to detect animals and their movement was automatically tracked. Movement parameters such as instantaneous speed, mean speed, and directionality (forward or reverse) were calculated from analyses of the tracks of individual animals. For calculating instantaneous speed, a moving average speed over 3 s was smoothed using locally weighted polynomial regression. Reverse movement was defined as movement of the head of the animal in the reverse direction at a minimum absolute speed of 30 µm/s that lasted for at least 1 s. The fraction of time spent reversing (Figure 4E, H) was calculated as the time spent reversing divided by the total time over which the animal was tracked. Turns (Figure 4F, I) were annotated manually and categorized as omega-turns, reversal-coupled omega turns, or delta turns using Fiji (Figure S4F) [75–77]. A turn was categorized as an omega turn if the animal assumed an Ω-like shape, where the head and the tail touched each other, or if the closest points near the head and the tail made an angle of <30 degrees relative to the deepest part of the mid-body (Figure S4F) [75, 76]. Delta turns were assigned if the animal assumed a δ-like shape and executed a deeper turn relative to an omega-turn by overturning onto itself as described previously [77]. Data were graphed using GraphPad Prism v9.3.1.

### QUANTIFICATION AND STATISTICAL ANALYSIS

Statistical tests were performed using GraphPad Prism v10.1.1. Specific statistical tests used are indicated in the figure legends. Normality was determined using a D’Agostino-Pearson omnibus normality test; if data were not normally distributed, non-parametric tests were used. Power analysis was performed with G*Power v3.1.9 [78]. Results of all statistical tests reported in the manuscript are shown in Dataset S1.

