## Supplemental Material for "Carbon dioxide shapes parasite-host interactions in a human-infective nematode"

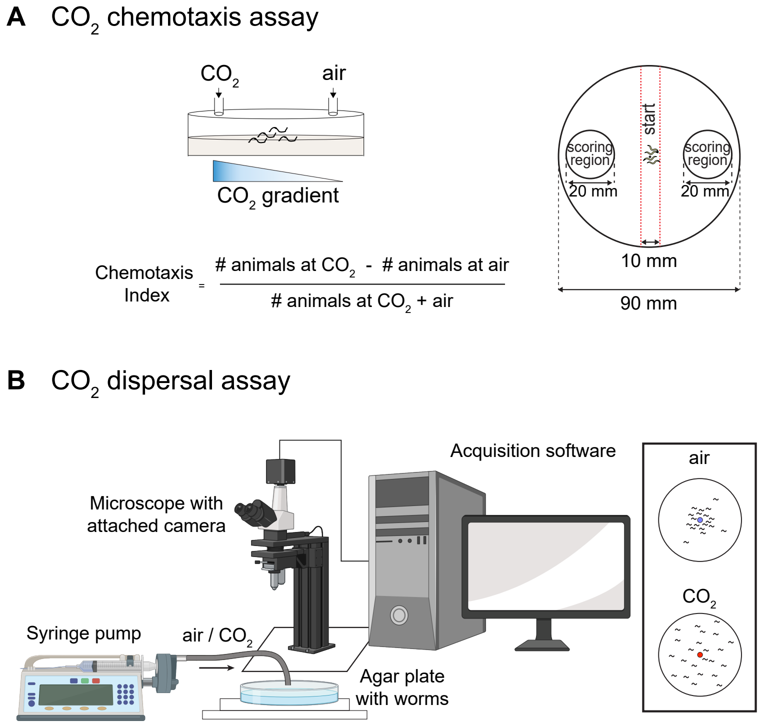


**Figure S1. $CO_2$ chemotaxis and dispersal assays. Related to Figure 1. A.** Schematic of the $CO_2$ chemotaxis assay for assessing the movement of parasitic nematodes in a $CO_2$ gradient [1, 2]. Left, side view of the assay plate; right; top-down view of the assay plate. Animals were placed along the center line of a 9 cm agar plate (right) and a $CO_2$ gradient was generated by pumping $CO_2$ into one side of the plate and air into the other side through holes in the plate lid (left) using a syringe pump. At the end of the assay, the number of animals in each scoring region was counted and used to calculate a chemotaxis index according to the formula shown. Diagrams adapted from Banerjee *et al.,* 2024 [3]; worms are not to scale. **B.** Schematic of the $CO_2$ dispersal assay for assessing dispersal of iL3s from either a $CO_2$ source or an air control source. Gases were pumped into the center of a 9 cm agar plate and images of the plate were acquired for 5 min at 2 frames/s using a microscope with an attached camera. The final distance of each iL3 from either the $CO_2$ source or the air control source was then calculated *post hoc*. Schematic generated using BioRender.

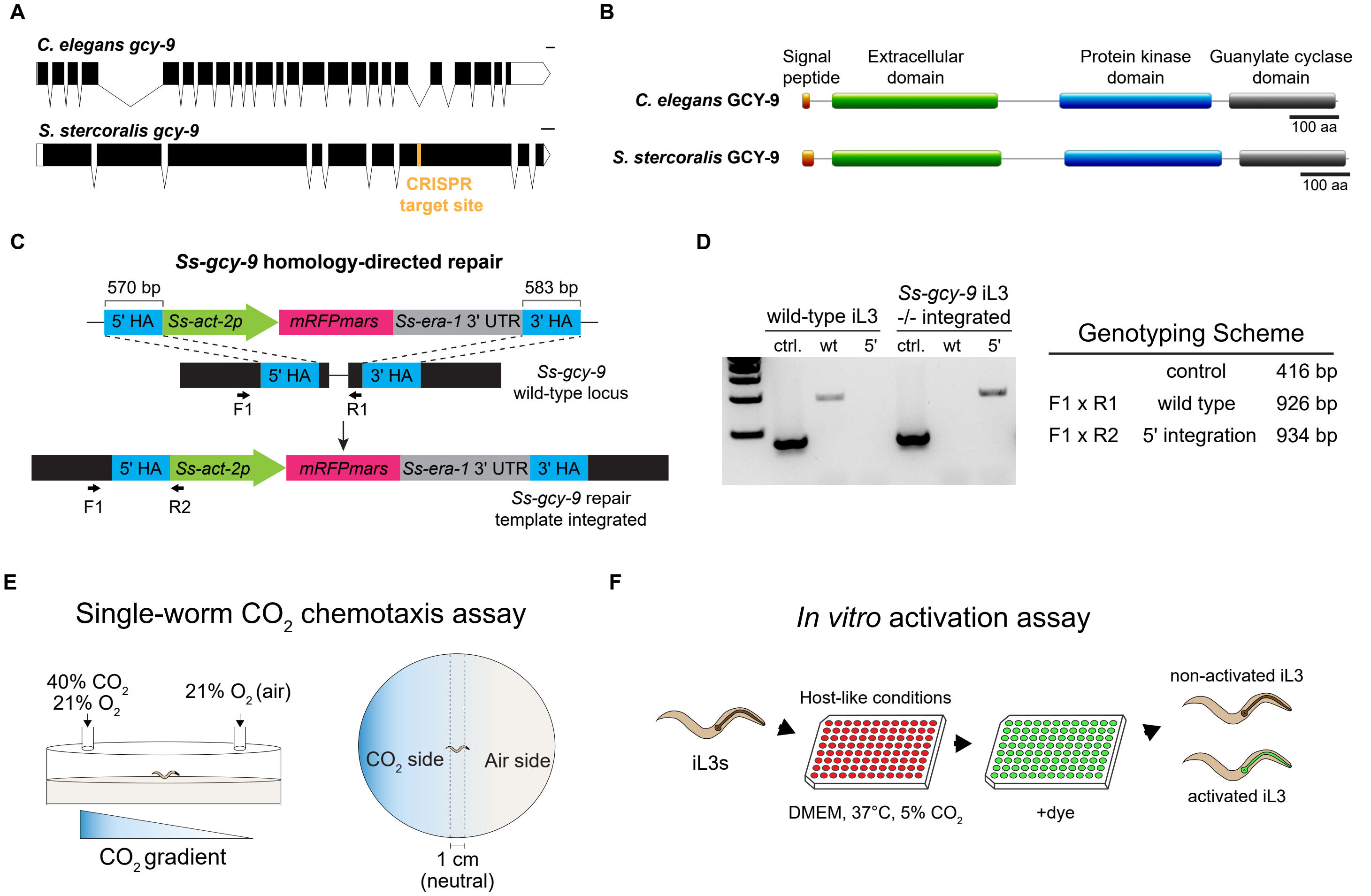


**Figure S2. Disruption of the *Ss-gcy-9* gene using CRISPR/Cas9-mediated targeted mutagenesis. Related to Figure 2. A.** Intron-exon diagram of the *gcy-9* genes of *C. elegans* and *S. stercoralis*. The *Ss-gcy-9* gene structure is based on the gene annotations from WormBase ParaSite [4, 5]. White bars indicate untranslated regions (UTRs); lines indicate introns. Scale bar = 100 bp. **B.** The predicted domains of the *C. elegans* and *S. stercoralis* GCY-9 proteins. The CRISPR target site is in the predicted guanylate cyclase domain of the *S. stercoralis* GCY-9 protein. Domain predictions are from InterPro and the figure was generated using ProSite. **C.** Strategy for CRISPR/Cas9-mediated disruption of the *Ss-gcy-9* gene using homology-directed repair. HA = homology arm. Successful integration of the repair template will insert *mRFPmars* into the *Ss-gcy-9* gene, thereby disrupting the gene. Expression of the mRFPmars reporter is driven by the promoter for the *Ss-act-2* gene [6]. The wild-type locus surrounding the *Ss-gcy-9* target site is amplified by the F1 x R1 primer set. Successful integration of the 5' HA of the repair template is confirmed by the F1 x R2 primer set. Primer binding sites shown are approximate. **D.** Gel showing representative genotyping of a wild-type and *Ss-gcy-9*$^{-/-}$ iL3 (left), and the corresponding genotyping scheme (right). iL3s with a homozygous disruption of the *Ss-gcy-9* gene and successful integration of the repair template will lack the wild-type (wt) band that amplifies a 926 bp region around the CRISPR target site and will have a 5' integration band (5') that amplifies a 934 bp region spanning the end of the 5' homology arm and part of the upstream genomic DNA. The control (ctrl) band amplifies a portion of the *Ss-act-2* gene and serves to confirm that genomic DNA is present [6]. The binding sites for the genotyping primers are shown in C. **E.** Schematic of a single-worm $CO_2$ chemotaxis assay. Left, side view of the assay plate; right, top view of the assay plate. A single iL3 is placed in the center of a 9 cm agar plate and a $CO_2$ gradient is established by pumping 40% $CO_2$ into one side of the plate and air into the other side through holes in the plate lid. At the end of the 10-min assay, the side of the plate containing the iL3 is determined. iL3s that remain in the center zone of the plate ("neutral," right) are not counted. **F.** Schematic of an *in vitro* activation assay [7-9]. iL3 are incubated in host-like conditions (DMEM, 37°C, 5% $CO_2$) for 21 h, after which fluorescent dye is added to the wells. Activated iL3s that resume feeding ingest the dye, resulting in pharyngeal fluorescence. Non-activated iL3s that have not resumed feeding do not have a fluorescent pharynx. Schematic is from Gang *et al.,* 2020 [7].

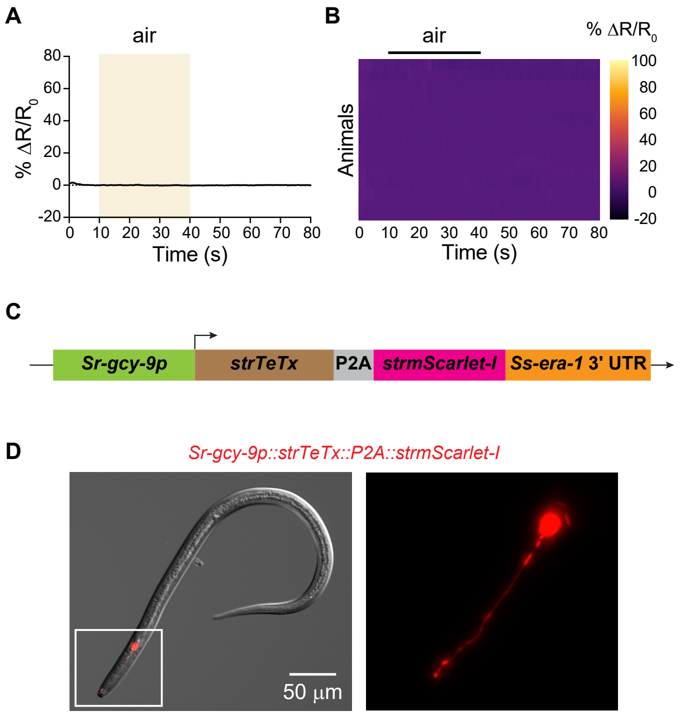


**E** ***Strongyloides*** **codon-optimized tetanus toxin (*strTeTx*) sequence**

atgccaattactattaataatttccgttattctgatccagttaataatgatactattattatgatgga
accaccatattgtaaaggacttgatatttattataaggctttcaaaattactgatcgtatttggatt
gttccagaacgttatgagttcggaactaaaccagaagatttcaatccaccatcttctcttattga
aggagcttctgaatattatgatccaaattatcttcgtactgattctgataaagatcgtttccttcaa
actatggttaaacttttcaatcgtattaagaataatgttgctggaggtaagtttaaacatatatat
actaactaaccctgattatttaaattttcagaagctcttcttgataagattattaatgctattccata
tcttggaaattcttattctcttcttgataagttcgatactaattctaattctgtttctttcaatcttcttga
acaagatccatctggagctactactaagtctgctatgcttactaatcttattattttcggaccagg
accagttcttaataagaatgaagttcgtggaattgttcttcgtgttgataacaaaaattatttccc
atgtcgtgatggattcggatctattatgcaaatggctttctgtccagaatacgttccaactttcga
taatgttattgaaaatattacttctcttactattggaaaatctaaatatttccaagatccagctcttc
ttcttatgcacgaacttattcatgttcttcatggactttacggaatgcaagtttcttctcatgaaatt
attccatctaaacaagaaatttatatgcaacatacttatccaatttctgctgaagaacttttcactt
tcggaggacaagatgctaatcttatttctattgatattaaaaatgatctttatgaaaaaactctta
atgattataaagctattgctaataaactttctcaagttacttcttgtaatgatccaaatattgatatt
gattcttataaacaaatttatcaacaaaaatatcaattcgataaagattctaatggacaatata
ttgttaatgaagataaattccaaattctttataattctattatgtatggattcactgaaattgaactt
ggaaaaaaattcaatattaaaactcgtctttcttatttctctatgaatcatgatccagttaaaattc
caaatcttcttgatgatactatttataatgatactgaaggattcaatattgaatctaaagatctta
aatctgaatataaaggacaaaatatgcgtgttaatactaatgctttccgtaatgttgatggatct
ggacttgtttctaaacttattggactttgtaaaaaaattattccaccaactaatattcgtgaaaat
ctttataatcgtactgctcgt

**Figure S3. BAG neurons respond to $CO_2$. Related to Figure 3. A.** *Ss*-BAG neurons do not respond to an air control. Graph shows the calcium response (mean ± SEM) of the *Ss*-BAG neurons to a 30 s pulse of air. Calcium response was measured using the ratiometric calcium indicator yellow cameleon YC3.60. Beige square shows the timing and duration of the air pulse. n = 15 iL3s. **B.** Heatmap of the *Ss*-BAG calcium responses. Each row shows the response of a single animal. Response magnitudes (% $\Delta R/R_0$) are color-coded according to the scale shown to the right. Rows are ordered by hierarchical cluster analysis. Black bar shows the timing and duration of the air pulse. **C.** Schematic of the construct used for cell-specific silencing of the *Ss*-BAG neurons. The *Sr-gcy-9* promoter was used to drive specific expression of tetanus toxin (TeTx) in the *Ss*-BAG neurons. The self-cleaving peptide P2A was used to express *strTeTx* and *strmScarlet-I* from the same promoter so that animals expressing TeTx could be identified based on expression of mScarlet. **D.** The TeTx transgene is expressed specifically in the *Ss*-BAG neurons. Left, representative DIC/epifluorescence overlay image of an *S. stercoralis* iL3 expressing TeTx in the *Ss*-BAG neurons. The head region indicated by the white box is enlarged to the right. Right, enlarged epifluorescence image of a BAG neuron; head is to the left. **E.** The synthesized cDNA sequence of the *Strongyloides*-codon-optimized *tetanus toxin* gene (*strTeTx*) that was used to silence the *Ss*-BAG neurons of *S. stercoralis* iL3s. The stop codon was removed for co-transcription with *strmScarlet-I* in the bicistronic vector shown in C. Purple font indicates a synthetic intron sequence.

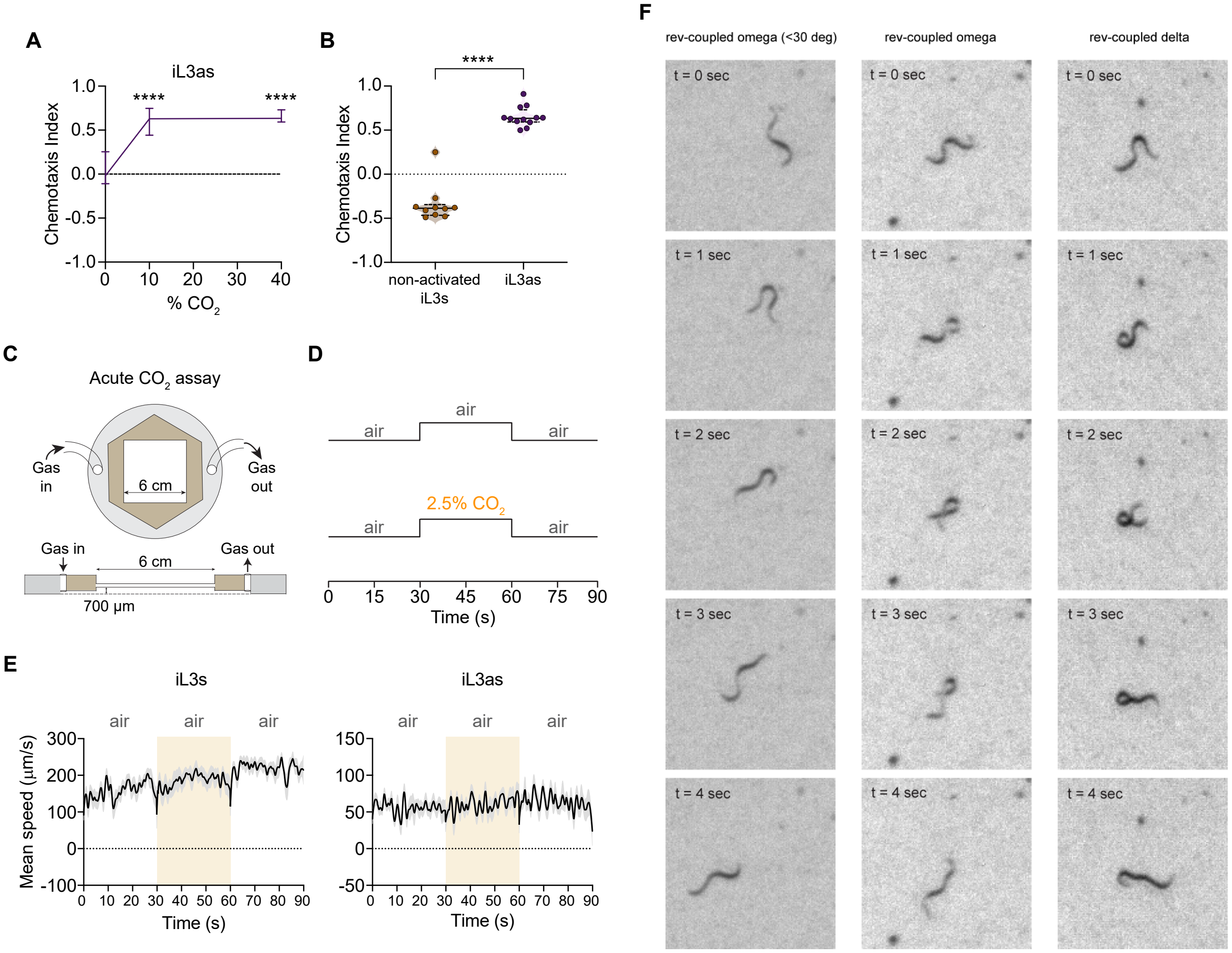


**Figure S4. iL3as are attracted to $CO_2$. Related to Figure 4. A.** iL3as are attracted to $CO_2$ concentrations as high as 40% in a $CO_2$ chemotaxis assay. Assays were conducted at 37°C to mimic intra-host temperature conditions. ****$p$<0.0001 relative to the 0% $CO_2$ control, two-way ANOVA with Dunnett's post-test. n = 12-16 trials per condition. Graph shows medians and interquartile ranges. **B.** $CO_2$ attraction is not the result of exposure to host-like conditions; rather, it is specific to iL3as. iL3s that were exposed to *in vitro* activation conditions but did not activate ("non-activated iL3s") were repelled by $CO_2$, while animals that activated under the same conditions were attracted to $CO_2$. Graph shows responses to 40% $CO_2$. Each data point represents a single $CO_2$ chemotaxis assay. Lines in violin plot indicate medians and dotted lines indicate interquartile ranges. ****$p$<0.0001, Mann-Whitney test. n = 10-12 trials per condition. **C.** Schematic of the chamber used to track the locomotion of iL3s and iL3as in response to acute $CO_2$ pulses. Animals are placed on an agar surface and covered by the gas-delivery chamber. Their locomotion is then video-recorded during gas exposure through the transparent 6 cm arena. Schematic is from Banerjee *et al.,* 2023 [10] and was adapted from Rojo Romanos *et al.,* 2018 [11]. **D.** The stimulus delivery paradigm for tracking responses to acute $CO_2$ pulses. In control assays, animals were exposed to 30 s of air, followed by 30 s of air from a second air tank, followed by 30 s of air from the original air tank. In experimental assays, animals were exposed to 30 s of air, followed by 30 s of 2.5% $CO_2$, followed by 30 s of air. **E.** Mean smoothed instantaneous speeds (± SEM) of animals from air control assays for iL3s (left) and iL3as (right). Responses of each life stage during the middle 30 s air pulse are quantified in Figure 4. n = 16-22 animals per condition. **F.** Representative time-series images of iL3s illustrating the different types of $CO_2$-evoked turns. Movements defined as turns were either reverse-coupled omega turns, where the head and tail made an angle of less than 30° (left column, t = 1 sec); reverse-coupled omega turns, where the head and tail touched (middle column, t = 2 sec); and reverse-coupled delta turns (right column, t = 1 sec) [12].

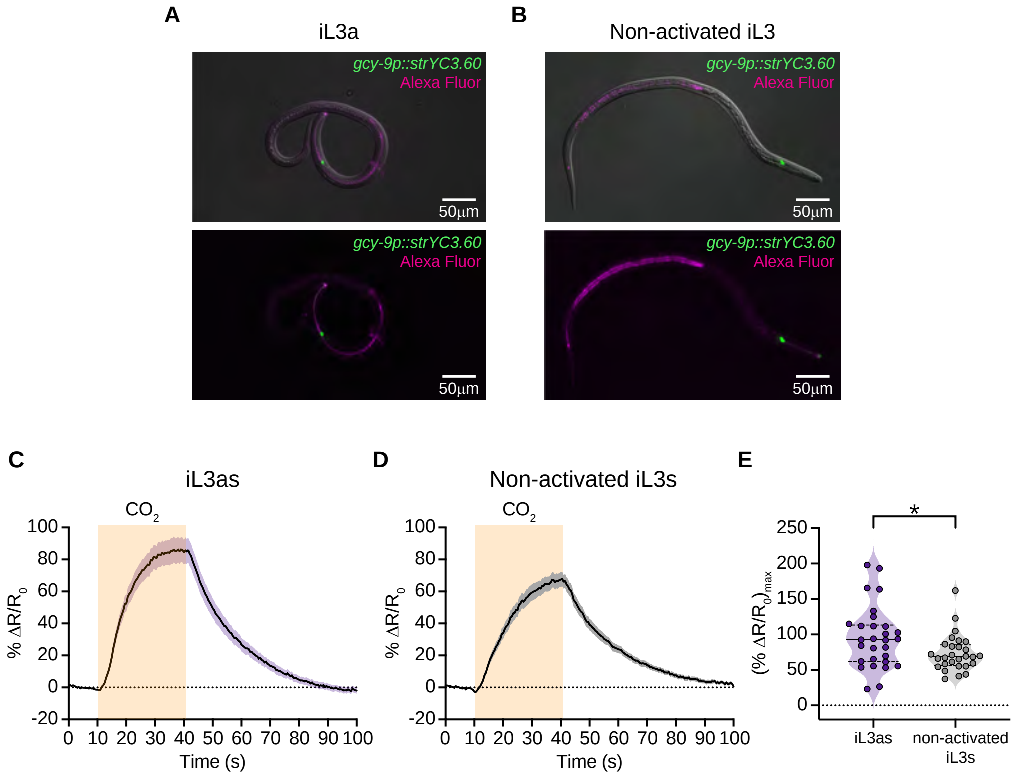


**Figure S5. iL3as show an enhanced $CO_2$-evoked calcium response in *Ss*-BAG neurons compared to non-activated iL3s. Related to Figure 5. A-B.** To perform calcium imaging from iL3as, an *in vitro* activation assay was first performed with iL3s expressing a *gcy-9p::strYC3.60* transgene. iL3as were identified based on the presence of Alexa Fluor NHS Ester 594 in the pharynx (A); the red Alexa Fluor dye was used instead of FITC so that the dye would not be visible during calcium imaging. iL3s that were exposed to the same activation assay conditions but did not activate ("non-activated iL3s") were identified by the lack of Alexa Fluor dye in the pharynx (B). The red fluorescence in the posterior part of the animal in B is due to intestinal autofluorescence. **C-D.** iL3as (C) show a larger $CO_2$-evoked calcium response in the *Ss*-BAG neurons than non-activated iL3s (D), as confirmed by statistical comparison of the maximum responses to the $CO_2$ stimulus (E). *$p<0.05$, Mann-Whitney test. n = 28-29 animals per life stage.

### Supplemental Tables

**Table S1. List of plasmids that were used in this study.** Plasmids are listed in the order in which they appear in the figures.

| Plasmid | Description |
|---|---|
| **pASB55**<br>*Sr-gcy-9p::strYC3.60::era-1* 3' UTR | Construct for specific expression of a *Strongyloides*-codon-optimized yellow cameleon *YC3.60* gene in BAG neurons |
| **pPV540**<br>*Sr-eef-1Ap::strCas9::Ss-era-1* 3' UTR | Construct for expression of *Strongyloides*-codon-optimized *Cas9* |
| **pSSG05**<br>*Sr-U6* 5' UTR*::Ss-gcy-9-sgRNA::Sr-U6* 3' UTR | Single guide RNA construct for CRISPR-mediated disruption of *Ss-gcy-9* |
| **pSSG04**<br>5' HA::*Ss-act-2p::mRFPmars*::*Ss-era-1* 3' UTR::3' HA | Homology-directed repair construct for *Ss-gcy-9* |
| **pNB11**<br>*Sr-gcy-9p::strTeTx::P2A::strmScarlet-I::Ss-era-1* 3' UTR | Construct for specific expression of a *Strongyloides*-codon-optimized *tetanus toxin* gene in BAG neurons |

**Table S2. List of primers that were used in this study.** Primers are listed in the order in which they appear in the figures.

| Primer | Description |
|---|---|
| **MLC168 ("*Sr-gcy9pro* HindIII F2")**<br>5'-ATA TAT AAG CTT TTG GAA GAA CAG CTC CTC TTT TAG TAA CAC-3' | 5' primer for amplification of the *Sr-gcy-9* promoter |
| **MLC169 ("*Sr-gcy9pro* BamHI R2")**<br>5'-ATA TAT GGA TCC AGA TGA ATC ACT TT T TGA AAT ATC TTT TGA TGG CA-3' | 3' primer for amplification of the *Sr-gcy-9* promoter |
| **SG78 ("*Ss-act-2* exon 1 F1")**<br>5'-GTA TTC CCT TCT ATT GTT GGA AGA CC-3' | Primer for amplifying exon 1 of the *Ss-act-2* gene |
| **SG80 ("*Ss-act-2* exon 1 R1")**<br>5'-CCT TCA TAG ATT GGT ACA GTG TGA G-3' | Primer for amplifying exon 1 of the *Ss-act-2* gene |
| **SG142 ("*Ss-gcy-9.1* F1")**<br>5'-GGA TAC TTT GTC AAC GTG GTT CAT T-3' | Primer for amplifying the *Ss-gcy-9* wild-type locus near the CRISPR target site<br>*-and-*<br>Primer for testing for genomic integration of the 5' homology arm |
| **SG143 ("*Ss-gcy-9.1* R1")**<br>5'-GAC TTC CAC CAC CTT TTG ATT TAA GA-3' | Primer for amplifying the *Ss-gcy-9* wild-type locus near the CRISPR target site |
| **MLC95 ("*Ss* mutscreen R2")**<br>5'-CGA GGT ACC TCT TTT CCA CAC TT-3' | Primer for testing for genomic integration of the 5' homology arm |

**Table S3. Summary of microinjections introducing the CRISPR constructs for targeted disruption of the *Ss-gcy-9* gene.** Table columns show, from left to right, the number of free-living adults injected ($P_0$), the number of $F_1$ iL3 screened, the number of transgenic iL3s collected, the number of transgenic iL3s genotyped, the number and percentage of genotyped iL3s with integration of the repair template, and the number and percentage of genotyped iL3s with a homozygous disruption of *Ss-gcy-9*. Percentages in the last two columns were calculated based on the number of red iL3s genotyped by PCR. The bottom row displays the totals from all experiments.

| # free-living adults injected | # iL3s screened | # red iL3s collected | # red iL3s genotyped | # repair-template integrated (%) | # *Ss-gcy-9* knockouts (%) |
|---|---|---|---|---|---|
| 46 | 1,380 | 61 | 25 | 19 (76%) | 7 (28%) |
| 25 | 397 | 28 | 13 | 11 (85%) | 0 (0%) |
| 25 | 103 | 12 | 12 | 7 (58%) | 1 (8%) |
| 22 | 782 | 24 | 24 | 22 (91%) | 14 (58%) |
| 23 | 300 | 14 | 11 | 10 (91%) | 5 (45%) |
| 22 | 1,126 | 39 | 24 | 24 (100%) | 10 (42%) |
| **163** | **4,087** | **178 (4.3%)** | **109** | **93 (85%)** | **37 (34%)** |
